# A Membrane-Disruptive Action of VBIT-4 Challenges Its Role as a Widely Used VDAC1 Oligomerization Inhibitor

**DOI:** 10.1101/2025.06.30.661942

**Authors:** Varun Ravishankar, Luís Borges-Araújo, Megha Rajendran, Elodie Lafargue, Deborah Byrne, Nicolas Buzhinsky, Mya S. Wolfe, Wendy Fitzgerald, Nina A. Bautista, Bethel G. Beyene, Motahareh G. Larimi, Jean-Pierre Duneau, James Sturgis, Sergey M. Bezrukov, Ignacio Casuso, Tatiana K. Rostovtseva, Lucie Bergdoll

## Abstract

Voltage-dependent anion channel (VDAC) is the most abundant protein of the mitochondrial outer membrane and a key regulator of metabolite exchange and mitochondrial physiology. Its oligomerization has been proposed to control processes such as mitochondrial DNA release and membrane remodeling, yet the underlying mechanisms remain poorly defined. VBIT-4 has been widely used over the last decade as a putative inhibitor of VDAC1 oligomerization, despite limited mechanistic validation. Here, using high-speed atomic force microscopy (AFM), we directly visualized VDAC1 assemblies in lipid membranes and examined the effect of VBIT-4. Unexpectedly, VBIT-4 induced membrane defects and permeabilization at micromolar concentrations, independently of VDAC1. Quantitative AFM analysis further shows that VBIT-4 does not alter VDAC1 cluster organization. Complementary electrophysiology, microscale thermophoresis, and coarse-grained molecular dynamics demonstrate that VBIT-4 partitions into lipid bilayers, increases membrane permeability, and destabilizes membrane structure, without detectable effects on VDAC1 channel properties or assemblies. Consistent with this mechanism, VBIT-4 induces VDAC1-independent cytotoxicity in HeLa cells at concentrations above 10 µM. Together, these results demonstrate that VBIT-4 does not act as a specific inhibitor of VDAC1 oligomerization but instead functions as a membrane-active compound. This work provides a revised framework for interpreting studies using VBIT-4 and highlights the importance of systematically assessing drug–membrane interactions when targeting membrane proteins.

## Introduction

The Voltage-Dependent Anion Channel (VDAC) is the most abundant protein of the mitochondrial outer membrane (MOM), accounting for over half of its protein content[1, 2], and the primary conduit for metabolite and ion exchange between mitochondria and the cytosol. Beyond transport, VDAC also regulates diverse mitochondrial functions, including apoptosis[3, 4], and calcium homeostasis[5–7]. Mammals express three isoforms (VDAC1, 2, and 3), which share their channel transport function but diverge in interaction partners and pathway specificity[8, 9].

VDAC is often recognized as a drug target because of its links to diverse diseases[10–12]. Despite its essential roles in physiology, lack of a defined catalytic site[12], and incomplete understanding of its functions in vivo complicates therapeutic development. One proposed strategy, attracting increased interest in recent years, is to target VDAC oligomerization, which has been implicated in essential physiological functions, including mitochondrial DNA release[13], lipid scrambling[14], mitochondrial organization[15], and interaction with MOM proteins[16] — processes central to mitochondrial dynamics and stress adaptation. However, rather than forming discrete oligomeric complexes (e.g. dimers, tetramers…), VDAC assembles into heterogeneous lipid–protein clusters [1, 2, 17, 18], whose organization is strongly dependent on membrane composition[17], challenging their selective targeting.

In 2016, Ben-Hail *et al.* introduced VBIT-4 as a small-molecule inhibitor of VDAC1 oligomerization[19]. Since then, it has been widely used to probe VDAC1 in various diseases, ranging from autoimmunity and neurodegeneration to metabolic disease models[13, 20–25]. Despite this widespread use, the mechanism of action of VBIT-4 remains poorly defined, and its effects on VDAC1 oligomerization have not been directly validated at the structural or biophysical level. Importantly, current conclusions regarding VDAC1 oligomerization and its modulation by small molecules rely largely on indirect assays, such as chemical cross-linking or apoptosis readouts, which do not resolve the native organization of VDAC in membranes.

High-speed atomic force microscopy (HS-AFM) has recently enabled real-time, nanometer-scale visualization of VDAC1 oligomerization[17], showing that lipid composition is a key regulator and that disruption of physiological lipid distribution destabilizes native-like assemblies. Using HS-AFM, we examined the effect of VBIT-4 on VDAC1 assemblies and found that it induces membrane permeabilization and bilayer defects even in the absence of VDAC1, contrary to expectations. Then, using complementary methods—including single-channel electrophysiology, microscale thermophoresis, and molecular dynamics—we confirmed that VBIT-4 partitions into membranes and perturbs bilayer properties independently of VDAC. Cellular assays corroborated these findings by showing that VBIT-4 induces cytotoxicity above 10 µM and reduces mitochondrial respiration and membrane potential even below 10 µM. These findings demonstrate that VBIT-4 acts by perturbing lipid bilayers rather than through detectable direct modulation of VDAC1, and highlight the importance of systematically assessing drug–membrane interactions when studying membrane proteins.

## Results

### VBIT-4 induces perturbations in VDAC1-containing membranes

HS-AFM provides label-free, real-time imaging of nanoscale membrane dynamics under near-physiological conditions, making it well-suited for assessing small molecule effects on VDAC1 oligomerization. We applied this approach to POPC:POPE:cholesterol membranes, reconstituted with or without VDAC1 and adsorbed on mica. Before VBIT-4 addition, VDAC1-containing membranes showed the characteristic “honeycomb” topography described in [17] (**Figure 1A and F**), with extended protein assemblies separated by lipid areas. Addition of 1 µM VBIT-4 produced small perforations a few nanometers wide and deep (**Figure 1B\***), indicating local bilayer destabilization while VDAC1 assemblies remained structurally intact (**Figure 1G**). Height profiles clearly distinguished VDAC1 from deeper VBIT-4–induced defects. At 10 µM, the number and size of defects increased (**Figure 1C\***), while the VDAC organization is not affected (**Figure 1C and H**). AFM images revealed a clear difference between organized VDAC1 pores, which remained intact, and the irregular defects formed in lipid regions by VBIT-4 (**Figure 1C, left**). Height profile analysis showed that VBIT-4-induced pores are wider (10-20 nm) and deeper (3-4 nm) than those of VDAC1 (5-7 nm wide, 0.8-2.5nm deep), confirming their distinct origin. Furthermore, height distribution analysis (**Figure 1D**) shows that VDAC1 containing membranes without VBIT-4 yield a narrow peak around 1nm, whereas the addition of VBIT-4 broadens the distribution to -4 nm at 1 µM and up to -5.5 nm at 10 µM, reflecting heterogeneous pore formation.

**Figure 1.**
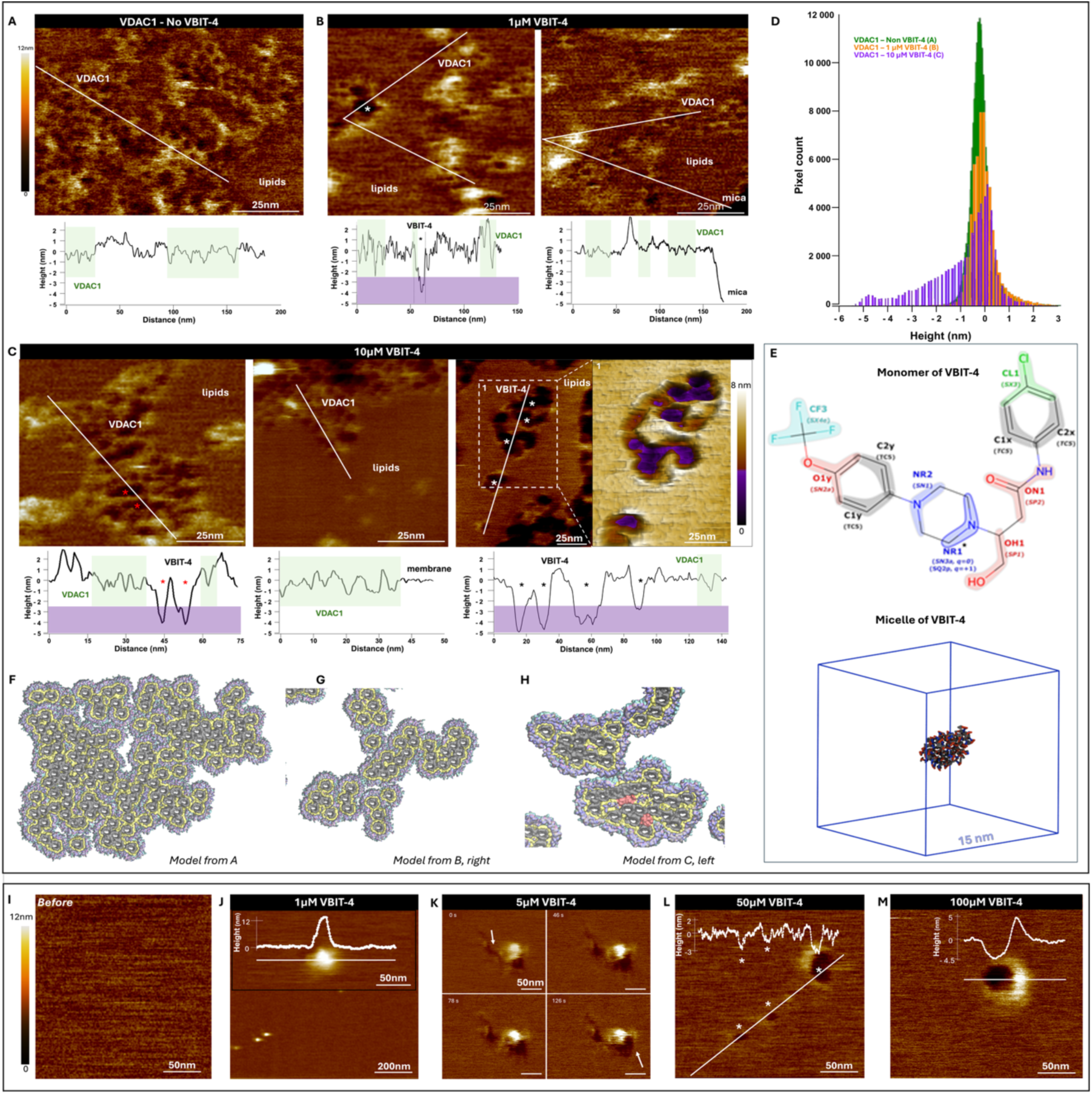
VBIT-4 induces defects in membrane bilayers, both in the presence and absence of VDAC1. **pper panels:** VDAC1 reconstituted into POPC:POPE:chol (60:32.5:7.5) liposomes was adsorbed onto mica for AFM imaging. Representative AFM images and corresponding height profiles before addition (**A**) and after addition of 1 µM (**B**) and 10 µM (**C**) VBIT-4. The purple rectangles in the height profiles indicate the 2.5 nm threshold: values above correspond to VDAC1 pores, whereas values below correspond to defects induced by VBIT-4. Asterisks (*) denote VBIT-4–induced pores. In (**C**), the left and middle panels show VDAC clusters, while the right panel presents an overview and a cross-section from a 3D zoom highlighting the depth of VBIT-4–induced pores in lipid regions devoid of VDAC1. Images acquired at 10 µM VBIT-4 display reduced resolution, consistent with compound-induced alterations of membrane properties. **D.** Quantitative pore size distribution derived from pixel depth/height analysis of the overview images shown in A, B, and C. Histograms represent the distributions without VBIT-4 (green, panel A), in the presence of 1 µM (orange, panel B) or 10 µM (purple, panel C) VBIT-4. **E.** Monomer and micellar structure of VBIT-4 obtained by molecular dynamic simulations . On the monomer structure, Polar and hydrogen-bonding groups are highlighted in red, ionizable/potentially charged groups in blue, neutral aromatic regions in grey, and strongly hydrophobic (lipophilic) substituents in green. Overall, the molecule displays an amphipathic organization, with a charged/polar core (blue/red) flanked by lipophilic aromatic groups (grey/green). **F. G. H.** Models based on AFM topography from panels A, B (right), and C (left), respectively. VDAC1 is shown as a dark solid surface, the first lipid shell in yellow, and surrounding lipids in atom color representation. The red patches in H. show the position of the VBIT-4 pores, which cannot be modeled by this method. **Lower panels:** Lipid membranes (without VDAC) were adsorbed onto mica and imaged by AFM. **I.** Control membrane before VBIT-4 addition. After addition of 1 µM (**J**), 5 µM (**K**), 50 µM (**L**), and 100 µM (**M**) VBIT-4. Height profiles are overlaid on the images. The false-colour scale is 12 nm in all panels.

To quantitatively assess whether VBIT-4 affects VDAC1 organization, we analyzed protein compaction within clusters using inter-protein distance measurements. Protein positions were extracted from AFM images to identify nearest neighbors and classified according to the approximate number of lipid layers between VDAC molecules. This analysis revealed no significant difference in VDAC organization between control, 1 µM, and 10 µM VBIT-4, indicating that VBIT-4 does not alter the spatial arrangement of VDAC or surrounding lipids within clusters (**Supplementary Figure 2**). Consistently, the absence of detectable structural changes in VDAC1 clusters (**Figure 1F–H**) supports a model in which VBIT-4 primarily perturbs the lipid matrix rather than protein assemblies.

### VBIT-4 induces membrane perturbation in protein-free lipid bilayers

To determine whether these effects depend on VDAC1, we next examined protein-free membranes using HS-AFM. Control bilayers displayed uniform topography without defects (**Figure 1I**). Addition of 1 µM VBIT-4 induced isolated protrusions (**Figure 1J**) up to tens of nanometers in diameter (**Figure 1J** inset), consistent with higher-order oligomers or aggregates. These protrusions likely reflect VBIT-4 aggregates, as coarse-grained molecular dynamics simulations show that VBIT-4 forms micelles rather than remaining monomeric (**Figure 1E**, Supplementary Figure 1). At 5 µM, deformations intensified, producing transmembrane defects (**Figure 1K**); at 50 µM— commonly used in cell studies—disruption was pronounced (**Figure 1L**), and at 100 µM HS-AFM revealed nanometer-scale perforations consistent with rupture (**Figure 1M**). Height profile analysis confirmed a progressive increase in defects dimensions: mean defect depth increased from 2.5-3 nm at 50 µM (**Figure 1L overlaid height profile**) to 4.5 nm at 100 µM VBIT-4 (**Figure 1M overlaid height profile**) while average defect diameters expanded from ∼20 nm to ∼50 nm over the same concentration range, demonstrating a dose-dependent destabilization of the lipid bilayer by VBIT-4. Thus, VBIT-4 alone progressively destabilizes membranes in a concentration-dependent manner, with detectable effects already at low micromolar concentrations (1–5 µM), independent of VDAC1, culminating in the formation of large pores at high concentrations. Notably, while these results demonstrate that membrane disruption by VBIT-4 is intrinsic and does not require VDAC1, defects appeared at lower concentrations in the presence of VDAC1, likely reflecting its lipid-scrambling activity[14], which may accelerate VBIT-4 transfer between leaflets and promote pore formation.

### VBIT-4 induces membrane defects

To confirm VBIT-4 induced membrane perturbation, we performed Laurdan fluorescence measurements in POPC:POPE:chol (62.5:30:7.5) liposomes. VBIT-4 induced a dose-dependent decrease in generalized polarization (GP), consistent with increased water penetration and lipid disorder (**Figure 2A**, **Supplementary Figure 3**). To directly assess membrane permeabilization, we performed a fluorescein leakage assay using large unilamellar liposomes of the same composition as in the Laurdan fluorescence experiments. At high internal concentrations, fluorescein self-quenches, whereas its release upon membrane disruption increases fluorescence due to dequenching. The addition of VBIT-4 triggered a gradual and concentration-dependent increase in fluorescence (**Figure 2B**), indicating progressive leakage consistent with membrane destabilization by the compound.

**Figure 2.**
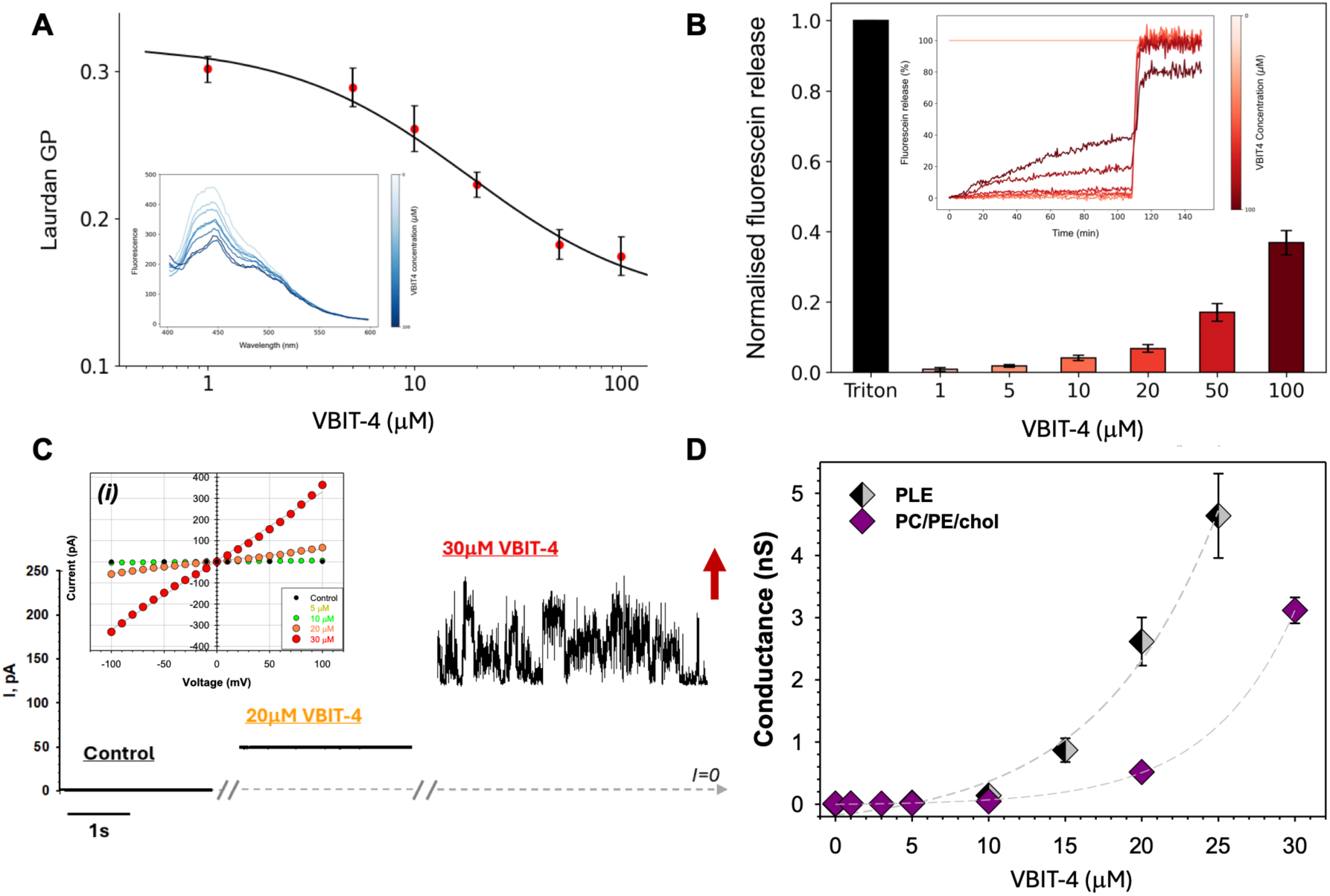
VBIT-4 incorporates into lipid membranes. **A.** Laurdan GP analysis of POPC:POPE:chol (62.5:30:7.5) liposomes with VBIT-4. The plot shows a VBIT-4 concentration-dependent decrease in Laurdan generalized polarization (n=5). The inset corresponds to the fluorescence spectra at VBIT-4 concentrations ranging from 0 (light blue) to 100 μM (dark blue); the inset data can be found at a larger scale in Supplementary Figure 4. **B.** Liposome leakage test of encapsulated fluorescein sodium salt from POPC:POPE:chol (62.5:30:7.5) liposomes with increasing concentration of VBIT-4 and TX-100 (0.5 %) (n=3). The inset represents the kinetics of fluorescein leakage from the liposomes. **C.** Representative current traces obtained on the same PLM made from DOPC:DOPE:chol (60:32.5:7.5) before and after consequent additions of 20 and 30 μM of VBIT-4 to the *cis* compartment of the chamber at 80 mV of applied voltage. Large fast fluctuations of the membrane conductance at 30 μM of VBIT-4 preceded the membrane rupture shown by the upward red arrow. The inset shows the current-voltage (I/V) curves obtained in the experiments at different VBIT-4 concentrations. Grey lines are linear regressions indicating the nearly Ohmic behaviour of VBIT-4-induced conductances. Dashed grey lines show a zero current. The membrane-bathing solutions consisted of 150 mM KCl buffered with 5 mM HEPES at pH 7.4. The current was digitally filtered using a 500 Hz Bessel (8-poles) filter for presentation. **D**. Conductance of planar membrane increases with VBIT-4 concentration. PLM were made from DOPC/DOPE/chol as in (C) and from PLE as in Supplementary Figure 4A. Membrane conductance was calculated from the corresponding I/V curves. The dashed lines are an exponential fit to guide the eye. Error bars are ±SD from measured conductances at different voltages.

To assess functional consequences of VBIT-4 insertion, we recorded currents across planar lipid membranes (PLMs) using the conventional voltage-clamp mode. VBIT-4 increased membrane conductance in a clear concentration-dependent manner, with rupture typically occurring between 20–50 µM across all lipid compositions tested—DOPC:DOPE:chol (62.5:30:7.5), soybean polar lipid extract (PLE), and DPhPC—despite experiment-to-experiment variability (**Figure 2C, D**; **Supplementary Figure 4A**; **Supplementary Table 1**). Representative traces illustrate progressive increase in conductance, reaching ∼3.5 nS at 30 µM VBIT-4 in DOPC:DOPE:chol and ∼5 nS at 25 µM VBIT-4 in PLE shortly before membrane rupture (**Figure 2C**, red arrow, **Figure 2D**; **Supplementary Figure 4A**). Large current fluctuations typically preceded PLM rupture, consistent with the formation of heterogeneous lipidic pores[26–28] rather than uniform channels. Similar behaviour has been reported for various amphiphilic compounds, which typically produce noisy current traces without well-defined discrete conductance levels[29–32]. VBIT-4–induced defects displayed nearly linear, symmetric current–voltage relationships (Inset in **Figure 2B** and **Supplementary Figure 4B**). These results demonstrate that VBIT-4 causes nonspecific membrane permeabilization above ∼15 µM. Notably, the onset of increased membrane conductance is observed at lower concentrations (∼5 µM), indicating that membrane perturbation precedes large-scale permeabilization. Our results differ from the original report[19], where 40 µM VBIT-4 did not permeabilize membranes formed by the ‘painted’ bilayer method using organic solvent. In contrast, when we repeated experiments with painted membranes (using Orbit mini system), VBIT-4 induced leakage starting from 5 µM (**Supplementary Figure 4C**), consistent with its effects in solvent-free PLMs. Together, these results demonstrate that VBIT-4 partitions into bilayers and disrupts their structure in a concentration-dependent manner, independently of VDAC, consistent with the AFM observations.

### VBIT-4 does not affect VDAC1 channel properties

Given the strong membrane perturbations observed above, we next tested whether VBIT-4 directly affects VDAC1 channel properties (conductance or voltage-gating). We performed single-and multichannel recordings at 5–30 µM VBIT-4. Across eight experiments, we observed no change in VDAC1 conductance or gating (**Figure 3A, B**). Instead, VBIT-4 induced a dose-dependent increase in overall membrane conductance (**Figure 3A**), leading to membrane rupture, which is consistent with the nonspecific leakage observed in protein-free membranes (**Figure 2C**). Multichannel recordings further confirmed that open probability (P_open)_ was unchanged (**Figure 3B**).

**Figure 3.**
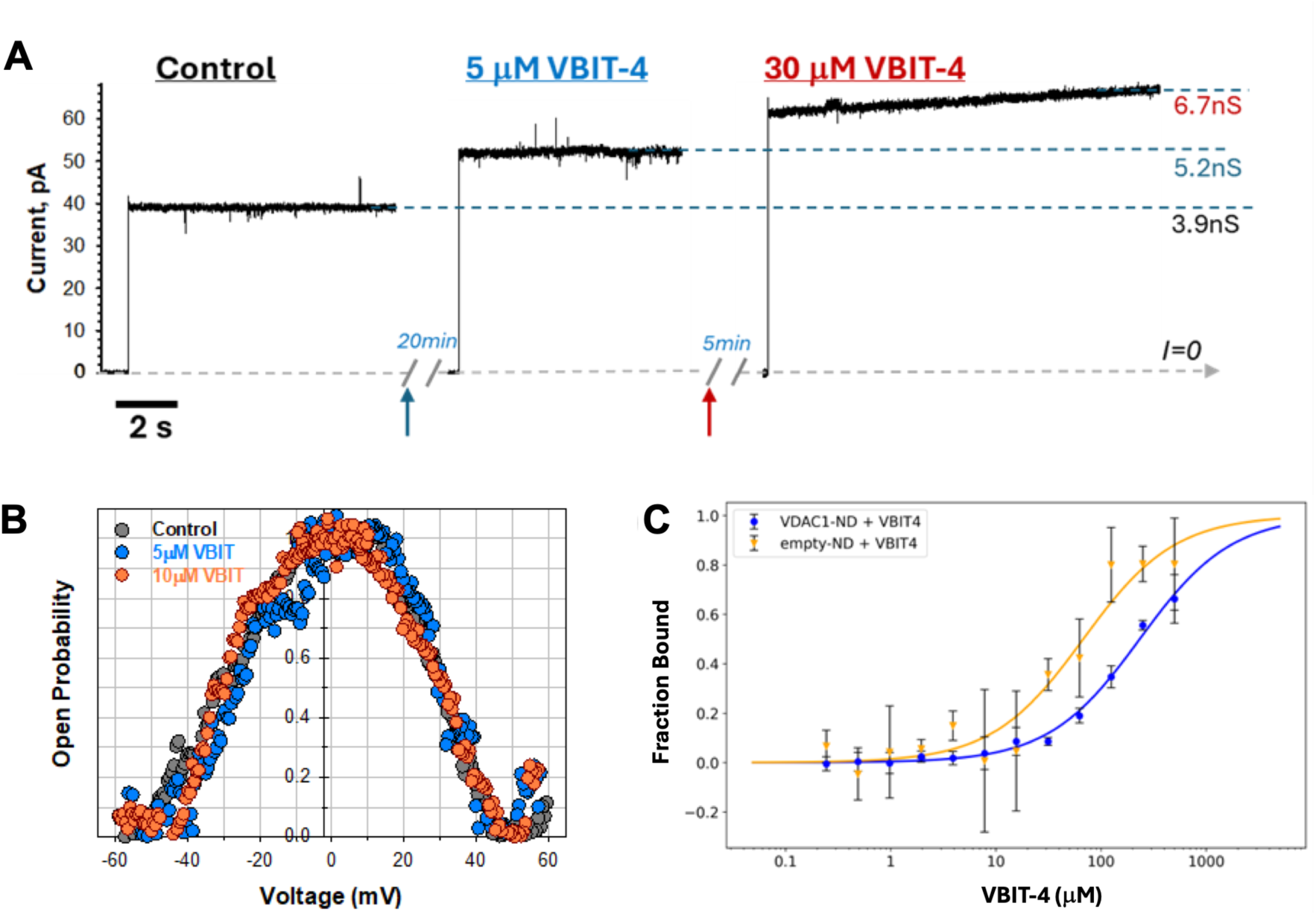
VBIT-4 destabilizes lipid membranes independently of VDAC1. **A.** Representative single-channel current traces obtained with VDAC1 reconstituted in PLM formed from PLE before (control) and after the addition of 5 and 30 μM VBIT-4 to the *cis* compartment (n=8). Addition of VBIT-4 and the time of the recordings after additions are indicated by upward arrows. The jumps in the current correspond to the application of 10 mV voltage following the application of 0 mV. The record was obtained on the membrane with the same single channel of 4 nS conductance as in the control. The grey dash line indicates zero current (*I=0*). The dashed blue lines indicate the current through the open single channel. Addition of 30 μM VBIT-4 induced a monotonic increase in membrane conductance, leading to membrane rupture. Current records were digitally filtered using an averaging time of 0.2 ms. **B.** Characteristic bell-shaped plots of open probability as a function of the applied voltage obtained in a multichannel experiment with VDAC1 in a PLE membrane in control and after addition of 5 and 10 μM VBIT-4. In all panels, the membrane-bathing solutions consisted of 1 M KCl buffered with 5 mM HEPES at pH 7.4. **C.** Binding of VBIT-4 to fluorescently labeled VDAC1 into nanodiscs (blue) or to empty nanodiscs, with MSP1D1 labeled (yellow), monitored by microscale thermophoresis. The plot shows normalized fluorescence at 650 nm measured 2.5 s after IR-laser activation across increasing VBIT-4 concentrations. An average apparent *K_D_* of 70 µM with a confidence interval from 20-167 µM (n = 4) was obtained for VBIT-4 binding to empty nanodiscs and 260 µM with a confidence interval from 105-331 µM (n = 2) for VDAC1-containing nanodiscs. Supplementary Figure 5B shows the control with MSP1D1 protein only.

To further probe VBIT-4’s interaction with VDAC1, we used Microscale thermophoresis (MST), a sensitive technique for detecting molecular interactions in solution. VBIT-4 titrated into lipid nanodiscs produced a Temperature-Related Intensity Change (TRIC) fluorescence signal (**Figure 3C**) with an average apparent dissociation constant (k_D)_ of ∼70 µM, comparable to reported affinities for VDAC1 in detergent micelles[19, 33]. A similar signal was observed for VDAC1-containing nanodiscs, but with a higher k_D o_f 260 µM, consistent with the reduced lipid content due to the space occupied by VDAC1. Together, these results indicate non-specific partitioning of VBIT-4 into the bilayer rather than direct interaction with VDAC1 (**Figure 3C**). Furthermore, no spectral shift in the 670 nm/650 nm fluorescence ratio was observed, indicating that the ligand does not induce changes in the immediate chemical environment of the target (**Supplementary Figure 5A**). These results indicate that VBIT-4 primarily partitions into lipid phases, with no detectable specific interaction with VDAC1. The variability across replicates, also observed in published titrations[19, 33], further supports membrane partitioning rather than binding to a defined protein site. The previously reported affinity likely reflects VBIT-4 partitioning into the hydrophobic core of detergent micelles[19, 33].

Together, these results demonstrate that VBIT-4 partitions into lipid bilayers and increases membrane permeability independently of VDAC1, without detectable direct interaction with or modulation of VDAC1 channel properties.

### Mechanistic insights into VBIT-4–mediated lipid bilayer destabilization from coarse-grained simulations

To provide a molecular interpretation of the experimentally observed membrane perturbations, we performed coarse-grained molecular dynamics (MD) simulations to explore how VBIT-4 partitions into and disrupts lipid bilayers independently of VDAC1. To reach longer timescales and larger system sizes, we used coarse-grained (CG) simulations with the Martini 3 force field[33], which reliably models membrane dynamics and small-molecule interactions, including with VDAC[34, 35]. This approach enables extensive sampling of membrane insertion and destabilization processes in a complex mimic of the MOM, that are not accessible at the all-atom level.

A CG model of VBIT-4 was parameterized from all-atom simulations (**Figures 4A and 4B**). Considering that VBIT-4 can exist in neutral and protonated states with distinct conformations, we developed separate models to analyze their effects.

**Figure 4.**
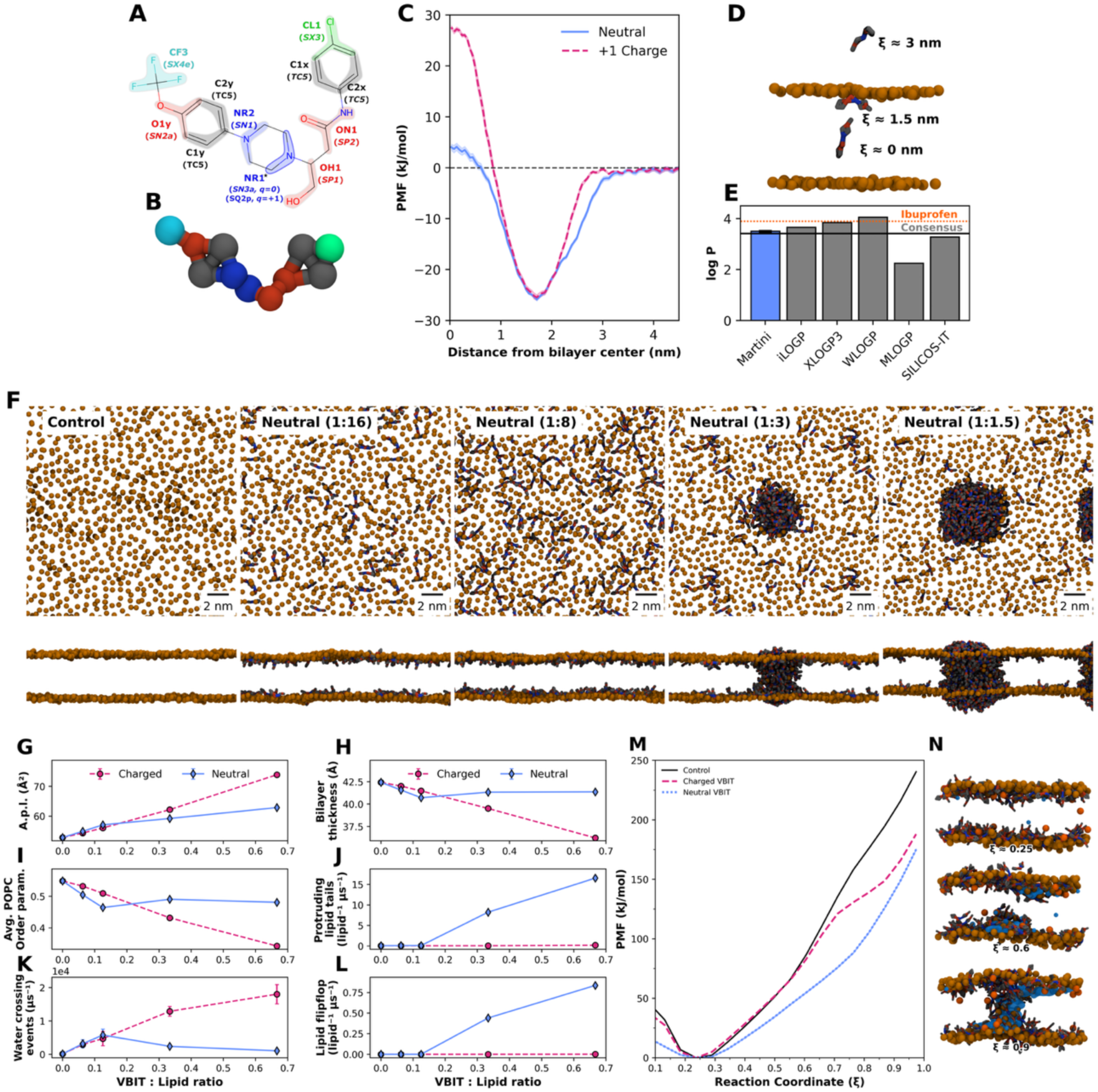
VBIT-4 partitions into lipid bilayers and disrupts membrane integrity through pore-like structures. **A.** VBIT-4 chemical structure and Martini 3 model mapping. Atom-to-bead mappings are indicated by the coloured shapes. Assigned Martini 3 bead names are indicated for each bead as overlaid bold text, with the corresponding bead types in italic text. Protonable group is highlighted (⋆). **B.** VBIT-4 Martini 3 CG model. **C.** Potential of Mean Force (PMF) of VBIT-4 insertion into coarse-grained MOM membranes. Free energy profiles are shown for both the neutral (blue) and protonated (+1 charge, red) forms of VBIT-4. **D.** Snapshots of the insertion of neutral VBIT-4 at different distances from the bilayer center (ξ); lipid PO_4 b_eads are represented in orange, and VBIT-4 in grey/red/blue. **E.** Predicted octanol–water partition coefficient (logP) values for VBIT-4. VBIT-4 hydrophobicity was estimated using several computational predictors (grey) and the Martini 3 coarse-grained model developed in this study (blue). Higher logP values indicate greater lipid solubility. The dashed black line shows the consensus value across predictors, while the orange dashed line marks the logP of ibuprofen (∼3.9), a well-known hydrophobic drug used here for comparison. **F.** Representative snapshots showing the impact of increasing VBIT-4 concentrations on MOM membrane mimics after 10 μs of CG MD simulation. (Lipid phosphate beads are shown in orange; VBIT in grey, with chemically distinct beads highlighted in red and blue). Top and side snapshots are shown for a 50:50 mix of charged and neutral VBIT-4 at increasing VBIT-4:lipid ratios (0, 1:16, 1:8, 1:3, 1:1.5). Lipid biophysical properties of MOM mimic membranes were evaluated in the presence of increasing VBIT-4 concentrations, either in the charged (red) or neutral (blue) form, as well as a 50:50 mixture of both states (orange). We monitored several indicators of membrane integrity: **G.** Area per lipid, **H.** Bilayer thickness, **I.** POPC acyl-chain order (membrane packing and organization), **J.** Lipid tail protrusion (packing defects), **K.** Bilayer water crossing (membrane permeability), and **L.** Lipid flip-flop (bilayer asymmetry). Each system was simulated for 10 µs in triplicate, and error bars represent the standard deviation across replicates. **M.** PMF profiles for polar defect formation in coarse-grained MOM membrane systems, shown as a function of the pore formation reaction coordinate ξ, for membranes without VBIT-4 (black), with charged VBIT-4 (red), and with neutral VBIT-4 (blue). Curve uncertainty (under the single-digit kJ mol^-1^ range) is represented by shading in the same colours along the corresponding curve. **N.** Representative snapshots of the membrane systems in the presence of neutral VBIT along the reaction coordinate; lipid PO_4 b_eads are represented in orange, cholesterol ROH in dark orange, water beads in blue (only those in the vicinity of the lipid headgroups or the established defect are shown), and VBIT-4 in grey.

### VBIT-4 readily inserts into lipid bilayers

To determine whether VBIT-4 can translocate across biological membranes—which is required to reach the mitochondrial membrane in cells—we performed umbrella sampling simulations to compute the Potential of Mean Force (PMF), which quantifies the free energy profile along the translocation pathway. As expected, the neutral form of VBIT-4 has a significantly lower translocation barrier (∼30 kJ/mol) compared to the charged form (∼55 kJ/mol, **Figure 4C**). While relatively high, these barriers are compatible with passive translocation, especially under conditions of elevated local concentration, membrane defects, transient pores, or lipid-scrambling activity, as suggested by our AFM experiments (**Figure 1**). It is important to note that the energy barrier estimated here applies to a single VBIT-4 molecule and does not account for cooperative or concentration-dependent effects. With a preferential position at around 1.5 nm from the bilayer core, VBIT-4 is positioned at the interface between polar heads and hydrophobic tails of lipids (**Figure 4D**). Overall, unbiased MD simulations show that VBIT-4 spontaneously inserts into lipid bilayers, consistent with its hydrophobic nature but also has the potential to translocate across it.

This membrane affinity is further supported by VBIT-4’s physicochemical properties. The octanol/water partition coefficient (logP_(oct/water))_ provides a measure of hydrophobicity: values below 0 indicating polar, water-soluble molecules (e.g., glucose, logP ∼ -3.2[36]), and values above 0 indicating apolar, lipophilic molecules (e.g., cholesterol, logP > ∼ 8[37]). Its predicted logP of ∼3.5 (**Figure 4E**), similar to ibuprofen (logP ∼ 3.9[38]), indicates strong membrane partitioning; a 1 µM aqueous concentration corresponds to ∼3.2 mM (10^3.5^=3,162.3) in the bilayer, highlighting its lipid preference.

In addition, simulations reveal a tendency of VBIT-4 to self-aggregate into micelle-like clusters, regardless of charge state (**Supplementary Figure 1**). These findings are consistent with Fourier-transform infrared spectroscopy (FTIR) measurements showing that VBIT-4 exists predominantly as aggregates, with only a minor soluble fraction (**Supplementary Figure 6**).

### VBIT-4 partitions into lipid membranes and induces membrane destabilization

We simulated a mitochondrial outer membrane mimic at increasing VBIT-4 concentrations (1:16– 1:1.5 VBIT-4:lipid). A recent predictor[39] estimates the pKa of VBIT4 to be between 5.7 and 6.9, consistent with the known piperidine pKa[40]. Because this range, close to physiological pH, allows coexistence of neutral and charged states, we simulated 50:50 mixtures, which produced pore-like aggregation and destabilization intermediate between the two extremes (**Figure 4F, G-L**). We also simulated both neutral and charged states of VBIT-4 independently (**Supplementary Figure 7**, **Figure 4G-L**). In all cases, VBIT-4 spontaneously inserted into the bilayer, increasing area per lipid, decreasing thickness and order, and enhancing water permeability (**Figure 4G-L**). At high concentrations, a neutral and 50:50 mixture of VBIT-4 aggregated to form pore-like structures that promoted lipid flip-flop and tail protrusion (**Figure 4G-L**). Charged VBIT-4 did not form stable pore-like structures but markedly increased permeability. Notably, the CG-Martini force field is known to over-stabilize membranes [41], suggesting that any observed destabilizing effects likely underestimate the actual impact on biological membranes. Despite this, the phase-separation and pore-like structures observed here are consistent with AFM and electrophysiology, supporting a mechanism in which VBIT-4 destabilizes bilayers at micromolar concentrations.

### VBIT-4 facilitates the formation of polar defects in lipid bilayers

To assess how VBIT-4 destabilizes membranes, we calculated the energy required to form polar defects—transient water-filled disruptions of the hydrophobic core. The energy required to form such defects determines membrane stability, with higher barriers preventing disruption and lower barriers making the membrane more susceptible to perturbation. To this end, we utilized the pore-formation reaction coordinate developed by Jochen Hub and applied umbrella sampling to probe this effect[42]. Control simulations without VBIT-4 yielded high barriers (∼250 kJ/mol) due to the known overstabilization of Martini 3, whereas all-atom DPPC simulations typically report ∼50 kJ/mol.

At ∼1:3 VBIT-4:lipid ratios, both charged and neutral VBIT-4 reduced the barrier by ∼60–100 kJ/mol (**Figure 4M, N**), indicating that the compound facilitates defect formation despite inflated absolute values in Martini. Although the small system size limited pore-like aggregation, unbiased simulations confirmed increased area per lipid, reduced thickness, enhanced water permeability, and frequent tail protrusions and flip-flop events—all consistent with reduced bilayer stability. Hence, even at low concentrations, VBIT-4 impacts the properties of biological membranes, independently of VDAC. These results are consistent with the AFM and electrophysiology data, supporting a membrane-driven mechanism.

### VBIT-4 Does Not Measurably Alter VDAC1 Cluster Organization

To directly assess whether VBIT-4 impacts VDAC1 oligomerization, we performed CG simulations of multiple VDAC1 molecules in the presence of increasing concentrations of VBIT-4 (**Supplementary Figure 8A,B**). In all conditions tested (VDAC1 alone and with VBIT-4 at VBIT:VDAC1 ratios of 20:1 and 55:1), systems consistently evolved toward similar oligomeric assemblies (**Supplementary Figure 8A,B**), with hyperbolic fits of the aggregation kinetics converging to statistically indistinguishable plateau values (**Supplementary Figure 8E,F**). This indicates that VBIT-4 does not measurably prevent or disrupt VDAC1 assembly under the simulated conditions.

At the highest concentration tested, a modest but statistically significant reduction in the extent of oligomerization was observed at 10 μs (p < 0.01), reflecting a concentration-dependent kinetic slowdown rather than a shift in the thermodynamic endpoint (**Supplementary Figure 8C, D**). This effect was only detectable at elevated VBIT-4 levels, approaching conditions where membrane defects begin to form (VBIT:VDAC1 55:1 ratio corresponds to a lipid:VBIT ratio of approximately 0.2) and was consistent across both protonation states. Importantly, oligomerization continued to progress throughout the trajectories, indicating that VBIT-4 does not alter the intrinsic propensity of VDAC1 clusters to assemble (**Supplementary Figure 8A,B**). Given the microsecond timescales accessible to these simulations, this transient kinetic hindrance is unlikely to be relevant to membrane organization on the timescales relevant in vitro or in cells.

While these simulations cannot address the behaviour of larger-order VDAC1 clusters likely present in vivo, the results are consistent with our AFM data, which support a model in which VBIT-4 has a negligible direct impact on VDAC1 clustering, with its primary effect arising from perturbation of the surrounding lipid environment. Notably, in the higher-concentration simulations, particularly with neutral VBIT-4, the VDAC1 oligomer appears to act as a nucleation point for VBIT-4 aggregation, with the resulting structures resembling the early stages of the pore-like membrane defects observed in our lipid-only simulations (**Supplementary Figure 8B**).

### VBIT-4 increases membrane permeability and cytotoxicity in HeLa cells

Given VBIT-4 membrane-disrupting effects, its partitioning into membranes could potentially contribute to cellular cytotoxicity. To assess whether these membrane effects translate to cellular systems, we assessed its cytotoxicity in live HeLa wild-type (WT) and VDAC1 knockout (KO) cell lines using the multitox-fluor multiplex cytotoxicity assay kit that reports on both membrane integrity and cell viability. In intact cells, the live-cell protease cleaves cell-permeant GF-AFC, leading to AFC fluorescence. Loss of membrane integrity leads to inactivation of live-cell protease and the release of dead-cell proteases, which can cleave the cell-impermeant bis-AAF-R110 and increase R110 fluorescence. This dual-measurement approach provides a comprehensive evaluation of the cytotoxic effects of VBIT-4, allowing us to directly correlate membrane disruption with cellular damage. There is a decrease in AFC fluorescence (viability) and an increase in R110 fluorescence (cytotoxicity) above 10 µM VBIT-4 with an IC_50 o_f ∼28-38 µM for HeLa WT (Viability: 32.59±1.04; Toxicity: 28.45±2.36) and VDAC1 KO (Viability: 37.43±2.66; Toxicity: 30.15±7.66) cell lines (**Figure 5A**). The observed VBIT-4 cytotoxicity is independent of VDAC1, corroborating our previous finding that VBIT-4 can perturb membranes and form pores above 10 µM, irrespective of VDAC1.

**Figure 5.**
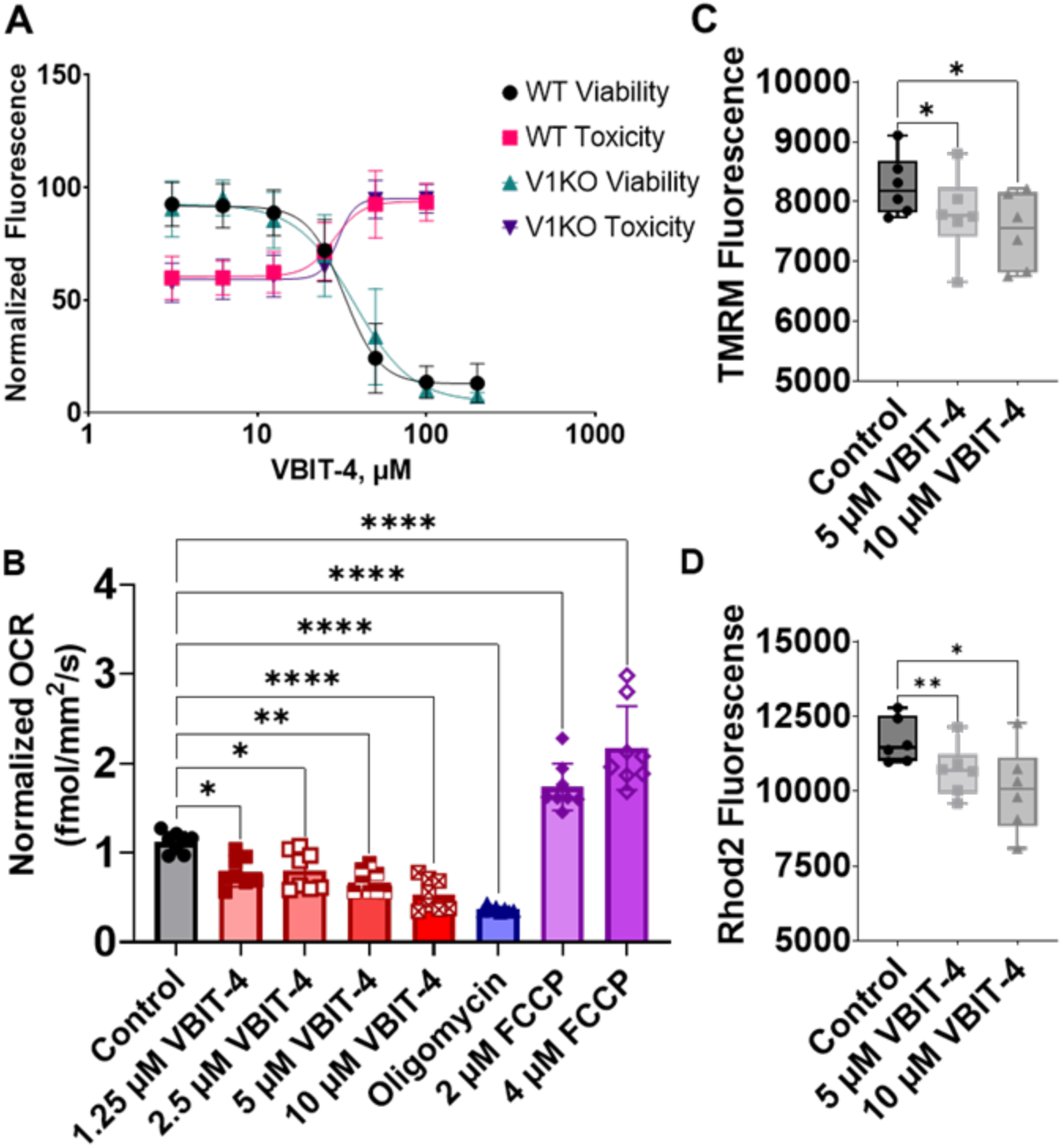
VBIT-4 decreases cell viability, mitochondrial respiration, and mitochondrial membrane potential in HeLa cells. **A.** The graph shows a decrease in viability (black and teal) and an increase in cytotoxicity (pink and purple) with increasing concentration of VBIT-4 in WT and VDAC1 KO HeLa cells. The data represent the average from 3 independent experiments, and the error bars represent the ±SD. The data was fit with nonlinear regression fit (GraphPad Prism 11.0.0). **B**. Bar graphs represent changes in the mitochondrial oxygen consumption rate (OCR) after 6 hours of treatment with 1.25 -10 µM VBIT-4 (red square symbols) compared to DMSO control (●) normalised to OCR before drug treatment. HeLa cells were treated with 0.5 µM oligomycin (▴), 2 µM (♦), and 4 µM FCCP (**◊**) as controls. Symbols represent 8 replicates, and error bars represent the ±SD from the mean. Flow cytometry measurements of mitochondrial membrane potential using TMRM (**C**) and mitochondrial calcium using Rhod2 **(D),** upon the addition of 5 µM (▪) and 10 µM (▴) VBIT-4 compared to vehicle control (●). Data from 6 independent experiments are represented in the box plots. The symbols represent data from independent experiments. The borders of the boxes define the 25^th^ and 75^th^ percentiles, with the median displayed as lines and error bars indicating the ±SD from the mean. Significance was tested using one-way ANOVA followed by the Dunnett post hoc test (*p < 0.05, **p < 0.01, ***p<0.001, ****p<0.0001).

We expect VBIT-4 to partition into multiple cellular membranes; however, because of the highly negative mitochondrial membrane potential, they are likely to attract the positively charged VBIT-4, prompting us to assess its effects on mitochondrial function. VBIT-4 reduced oxygen consumption rate (OCR) in a concentration-dependent manner (**Figure 5B**) within 6 hours of treatment. VBIT-4 also decreases the mitochondrial membrane potential measured with TMRM **(Figure 5C)**, without affecting mitochondrial mass (MTG) (**Supplementary Figure 9A**), and decreases mitochondrial calcium measured with Rhod-2 dye (**Figure 5D)**, as previously reported[43, 44]. Normalization of TMRM to MTG further confirmed a significant loss of mitochondrial potential at 10 µM VBIT-4 (**Supplementary Figure 9B**). These results could be explained by the membrane-destabilization activity observed in our in vitro experiments with VBIT-4.

Notably, the efficacy of VBIT-4 appeared to be dependent on storage conditions. Long-term storage at -20 °C results in complete loss of activity within two months, and even storage at -80 °C for more than a month leads to reduced efficacy compared to freshly prepared solutions (**Figure 6**). VBIT-4 is described by different manufacturers as an unstable molecule (one month at -20 °C, see the storage and stability information in selleckchem.com or other manufacturers). This could explain the wide variability in VBIT-4 concentrations used in various publications: the fresh samples are more cytotoxic, then this unstable molecule is modified. Together, these results from different techniques and systems suggest that VBIT-4 acts through membrane disruption rather than by directly modulating VDAC1.

**Figure 6.**
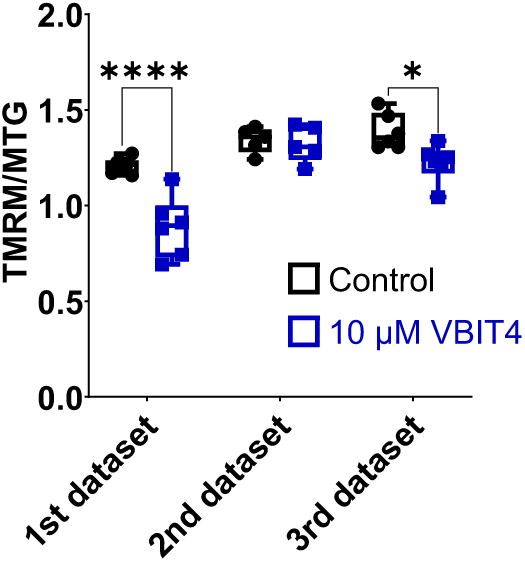
Storage conditions affect the cellular activity of VBIT-4. TMRM/MTG ratio comparing control (●) and 10 µM VBIT-4 (▪) stored in different conditions. 1^st^ dataset was collected within 2 weeks of preparation of fresh stock solution in DMSO. 2^nd^ dataset was using the same sample stored at -20 °C for 2 months, and the 3^rd^ dataset was using a sample stored at -80 °C for less than 1 month. Data from 5-6 independent experiments are represented in the box plots. The symbols represent data from independent experiments. The borders of the boxes define the 25^th^ and 75^th^ percentiles, with the median displayed as black lines and error bars indicating the ±SD from the mean. Significance was tested using one-way ANOVA followed by the Dunnett post hoc test (*p < 0.05, ****p<0.0001).

## Discussion

VBIT-4 was introduced in 2016 as a selective inhibitor of VDAC1 oligomerization, proposed to directly bind VDAC1 and prevent apoptosis triggered by cisplatin or selenite[19]. Since then, it has been widely adopted as a probe for VDAC1-dependent pathways in diverse disease contexts, including neurodegeneration, autoimmunity, and cancer[13, 45–48]. However, its mechanism of action was not well understood. Using a combination of biophysical methods and in vitro approaches, our results fundamentally challenge the interpretation of the assumed mechanism of action of VBIT-4. Across a suite of complementary approaches—HS-AFM, microscale thermophoresis, electrophysiology, Laurdan fluorescence, liposome-leakage assays, and MD simulations—we consistently observed that VBIT-4 partitions into lipid bilayers and destabilizes them. VBIT-4 inserts into the lipid bilayer, leading to lipid flip-flop, increased area per lipid, reduced bilayer thickness, and the formation of water-permeable defects. At micromolar concentrations (1–10 µM), these perturbations are already detectable and become more pronounced with increasing concentration of VBIT-4, in different lipid environments, independently of VDAC1, as shown by three independent techniques (AFM, electrophysiology, and liposome-leakage assay). Importantly, quantitative analysis of AFM images shows that VBIT-4 does not alter VDAC1 cluster compaction, as inter-protein distance distributions remain unchanged by the addition of VBIT-4 (**Supplementary Figure 2**). This indicates that, despite strong membrane perturbations, the organization of VDAC assemblies is preserved. Consistent with its membrane-disruptive activity, VBIT-4 exerts cytotoxicity in the micromolar range as previously reported[44]. We observed increased cell death in WT and VDAC1 KO HeLa cells, demonstrating that VBIT-4 toxicity is independent of VDAC1. The increased dead-cell protease activity, measured using a cell-impermeant fluorophore, suggests membrane permeabilization at concentrations greater than 10 µM VBIT-4. These findings demonstrate that VBIT-4 does not act as a specific VDAC1 inhibitor but rather functions as a membrane-active compound whose effects derive from general lipid membrane disruption, potentially affecting varieties of membrane proteins (**Figure 7**). Importantly, VBIT-4 was never shown to selectively target MOM in cells. By altering lipid organization, VBIT-4 potentially perturbs various cellular membranes and may influence numerous pathways beyond VDAC1, consistent with reports showing that it perturbs respiratory complexes in the inner mitochondrial membrane[44].

**Figure 7.**
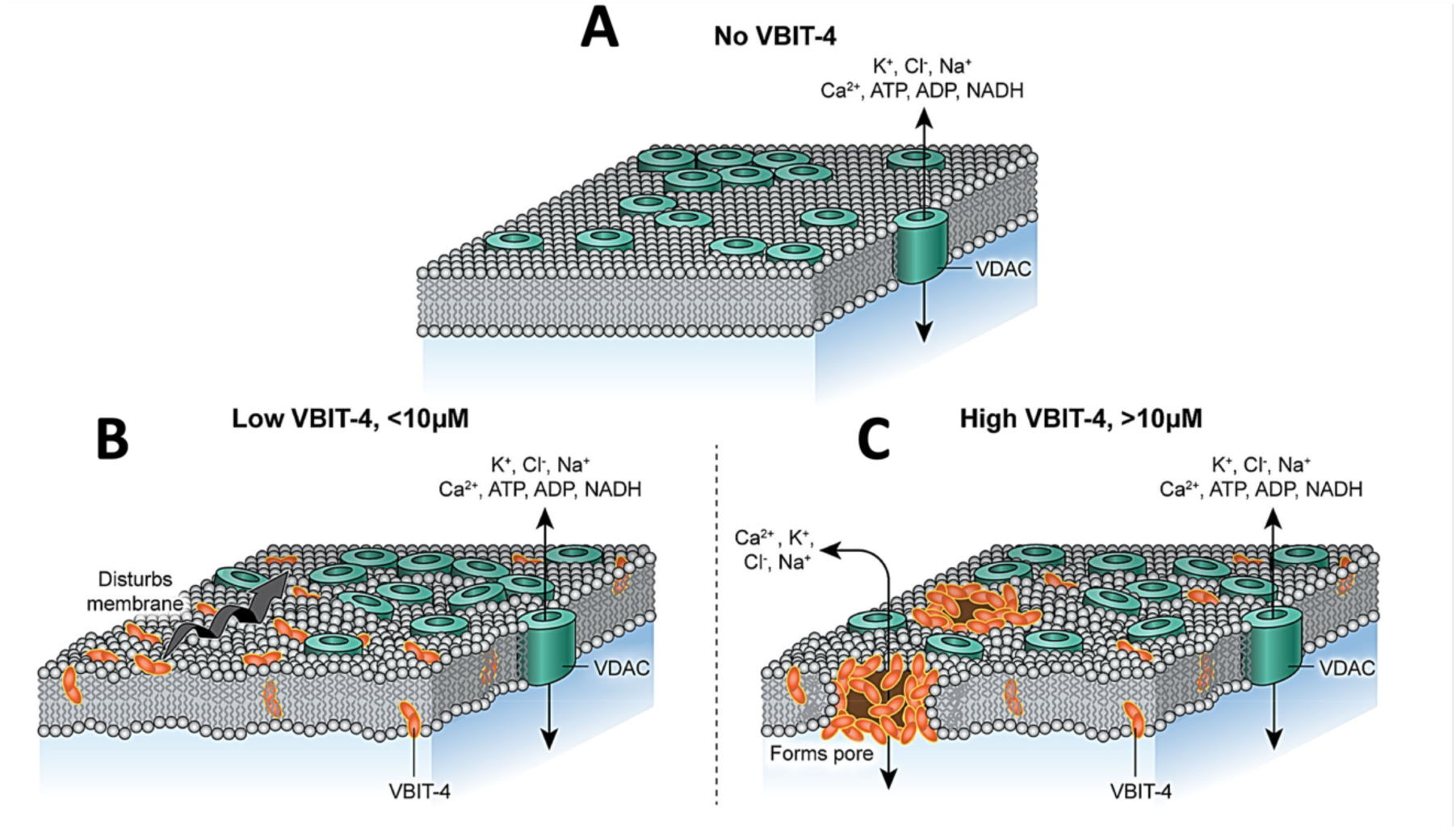
Putative model of VBIT-4 partitioning and pore formation into a VDAC1-containing membrane. **A.** Without VBIT-4, VDAC1 forms clusters of different sizes and compactions with both direct and lipid-mediated contacts between VDAC1 β-barrels. **B**. At low concentrations (< 10 µM), VBIT-4 partitions into the membrane, destabilizing it, without measurably altering VDAC1 cluster compaction. **C**. At high concentrations (>10 µM), VBIT-4 forms water-soluble pores in the membrane, inducing membrane permeabilization, further affecting membrane integrity. In all three scenarios, VDAC1 monomers maintain their transport channel properties, conducting the water-soluble metabolites, such as ATP, ADP, and NADH, and small ions, including calcium.

At concentrations below 10 μM, VBIT-4 decreased mitochondrial calcium, respiration, and membrane potential in HeLa cells without affecting mitochondrial mass. These findings align with reports that VBIT-4 also accumulates in the mitochondrial inner membrane, where it inhibits respiratory complexes I, III, and IV and decreases mitochondrial membrane potential[43, 44]. Notably, Belosludtsev *et al*. observed a decrease in calcium retention capacity at 30 μM VBIT-4 in their experiments on isolated mitochondria, which may indicate that VBIT-4 is a mitochondrial pore inducer. However, VBIT-4 also reduced the rate of calcium-dependent organelle swelling. This discrepancy was attributed to depolarization of the mitochondrial inner membrane, which inhibits both mitochondrial calcium retention and mitochondrial swelling[43].

All the above raise concerns regarding the inconsistencies in published cell data where VBIT-4 was used: while some studies report clear toxicity above 10–20 µM (our data and [44]), others describe apparent protection in pathological settings[13, 19, 49]. The variability can be attributed to various factors. First, the behaviour of VBIT-4 is strongly influenced by its poor chemical stability— consistent with our observation that mitochondrial effect disappears after only two months of storage in DMSO at −20 °C (**Figure 6**). Second, with two ionizable nitrogens in the piperazine ring, VBIT-4 exists at physiological pH as neutral and monocationic species that both display poor solubility, aggregation, and strong membrane partitioning. As a consequence, the nominal concentration of VBIT-4 used in experiments is unlikely to correspond to the effective concentration of active compound available in solution or within lipid membranes. The effective concentration is expected to vary substantially with pH, storage conditions, formulation, and membrane composition, complicating direct comparisons between different assays and across studies. Third, most studies use VBIT-4 at ∼10 µM—below overt cytotoxicity yet within the range where membrane perturbation occurs—and typically assess its effects in combination with apoptosis inducers or in disease models rather than measuring direct toxicity. Such discrepancies likely reflect differences in assay sensitivity, compound handling, and pH, as well as the altered membrane composition of diseased cells. In this context, “protective” effects are more plausibly attributed to nonspecific modulation of cellular membrane (including mitochondrial membrane) properties than to inhibition of VDAC1 oligomerization.

VDAC1 oligomerization has been proposed to regulate mitochondrial DNA release[13], lipid scrambling[14], and apoptosis[50], making it an attractive but controversial therapeutic target, as the implications of blocking VDAC1 oligomerization remain poorly understood. In addition, the structural basis of these assemblies remains elusive: except for a few defined interfaces[51–53], VDAC1 does not form rigid, stoichiometric oligomers but rather heterogeneous lipid–protein clusters[1, 2, 17, 18] whose size and compaction are strongly lipid-dependent[17]. Cross-linking— the assay most commonly used to monitor VDAC1 “oligomerization”— does not report on oligomer number or stability but rather on the proximity of proteins within these adaptable clusters in MOM. In contrast, HS-AFM directly measures the spatial organization of VDAC1 assemblies in lipid membranes, while molecular dynamics simulations provide an independent description of assembly behaviour. These approaches therefore probe related, but non-equivalent, aspects of VDAC1 organization. The organization of VDAC1 in the MOM is extremely sensitive to lipid composition[17]; any hydrophobic compound that perturbs membrane properties may therefore influence cross-linking efficiency through changes in protein spacing, orientation, or residue accessibility, without necessarily altering the overall organization of VDAC1 assemblies. Accordingly, although our results do not directly address cross-linking efficiency, AFM quantification (**Supplementary Figure 2**) and molecular dynamics simulations (**Supplementary Figure 8**) consistently show that VBIT-4 does not measurably alter VDAC1 cluster compaction or prevent assembly formation under the conditions examined.

Membrane perturbations could also affect VDAC’s interaction with calcium channels in other organelles such as endoplasmic reticulum, sarcoplasmic reticulum and lysosomes, thereby decreasing mitochondrial calcium uptake. Calcium overload is known to lead to apoptosis. Thus, the decrease in mitochondrial calcium could be linked to the protective effect of VBIT-4 at lower concentrations in pathological models. In conclusion, the prevailing view of VBIT-4 as a specific VDAC1 inhibitor requires re-evaluation, as its cellular activity is more consistent with nonspecific modulation of the lipid environment.

The nonspecific membrane activity of VBIT-4 highlights a broader challenge in drug discovery for membrane proteins. Hydrophobic small molecules frequently partition into bilayers, where they can alter lipid organization. Well-documented cases include licofelone, a COX/5-LOX inhibitor whose clinical development failed partly due to off-target membrane disruption[55], and omeprazole, which intercalates into DPPC bilayers and alters lipid domain structure[56].

Taken together, these findings demonstrate that VBIT-4 is not a suitable molecule to modulate VDAC1 oligomerization. Our results, therefore, caution against interpreting VBIT-4–induced phenotypes as evidence of VDAC1 inhibition. Importantly, these membrane effects occur within the same concentration range as those used in most studies invoking VBIT-4 as a specific VDAC1 inhibitor. Systematic assessment of drug–membrane interactions early in drug development is therefore essential, as the absence of rigorous lipid-only controls can result in membrane-driven effects being erroneously attributed to specific protein inhibition.

## Materials and Methods

### VBIT-4

VBIT-4 was purchased from Selleck Chemicals (VBIT-4, Cat. No.S3544, Selleckchem, Houston, TX, dissolved in DMSO for biophysical experiments, in powder for cellular assays) and stored at−80 °C according to the manufacturer’s instructions. For biophysical assays, aliquots were used within one year of purchase. For cellular assays, VBIT-4 was dissolved in DMSO, and aliquots were stored at −80 °C. Experiments were performed within 2 weeks of preparing the fresh solution, and aliquots were discarded after 3 freeze-thaw cycles.

### mVDAC1 purification

mVDAC1 production and purification were performed as previously described[57]. A pQE9 plasmid encoding mouse VDAC1 with an N-terminal 6×His tag was expressed as inclusion bodies in *Escherichia coli* M15 (pREP4) cells (Creative Biolabs). Cells were grown at 37 °C with agitation until reaching an OD₆₀₀ of 0.7, and protein expression was induced for 4 h with 1 mM IPTG. Cells were harvested by centrifugation and resuspended in 50 mM Tris-HCl pH 8.0, 2 mM EDTA, and 20% (w/v) sucrose. Lysis was performed by lysozyme treatment followed by sonication. Inclusion bodies were isolated by centrifugation for 15 min at 12,000 × *g*, washed with 20 mM Tris-HCl pH 8.0, 300 mM NaCl, and 2 mM CaCl₂, and solubilized in 20 mM Tris-HCl pH 8.0, 300 mM NaCl, and 8 M urea for 1 h. Insoluble material was removed by centrifugation at 25,000 × *g* for 1 h.

The supernatant containing solubilized mVDAC1 was loaded onto a Ni–NTA affinity column and eluted with 20 mM Tris-HCl pH 8.0, 300 mM NaCl, 8 M urea, and 150 mM imidazole. The purified protein was refolded by rapid dilution into 20 mM Tris-HCl pH 8.0, 150 mM NaCl, 10 mM DTT, and 1.5% (w/v) LDAO. Refolded protein was ultracentrifuged at 355,000 × *g* for 45 min to remove aggregates, concentrated, and further purified by size-exclusion chromatography on a Superdex 200 Increase 10/300 GL column (Fig. S11). The monomeric VDAC peak was collected, concentrated to 2 mg/mL, flash-frozen in liquid nitrogen, and stored in 20% (v/v) glycerol at −80 °C. Samples were ultracentrifuged after thawing prior to liposome reconstitution to remove residual aggregates.

### Reconstitution into liposomes

POPE, POPC, and cholesterol (Avanti Polar Lipids) were mixed in chloroform at 62.5:30:7.5 molar ratios and dried under a nitrogen stream. The lipid film was resuspended in 150 mM KCl, 20 mM Tris (pH 8.0) to a final lipid concentration of 12.5 mg/mL and vortexed thoroughly. The suspension was incubated at 37 °C for 10 min and sonicated for 1 min, or until small unilamellar vesicles (∼100 nm) formed, as verified by dynamic light scattering (DLS). For the VDAC1-containing liposomes, β-Octyl glucoside (β-OG) was added at a detergent-to-lipid ratio of 0.65 (w/w), followed by the addition of protein at a lipid-to-VDAC ratio of 3:1 (w/w). Detergent removal was performed overnight using Bio-Beads (Bio-Beads-to-detergent ratio 30:1 w/w). The resulting proteoliposomes were analyzed by DLS after reconstitution.

### AFM and HS-AFM data acquisition

To prepare protein-containing supported lipid bilayers (SLBs), 15 µL of proteoliposome suspension (1 mg/mL) was mixed with 85 µL of incubation buffer (10 mM HEPES pH 7.4, 100 mM NaCl, 15 mM CaCl₂) and incubated for 1 h at room temperature on freshly cleaved mica under a humid hood. The samples were rinsed ten times with ultrapure water (Milli-Q) and imaged in imaging buffer (10 mM HEPES pH 7.4, 100 mM NaCl, 5 mM CaCl₂) at 35 °C to verify membrane homogeneity. All conditions were tested in triplicate.

Conventional AFM imaging was performed in contact mode using a Nanoscope IIIe Multimode AFM (Bruker, Santa Barbara, CA, USA) equipped with OTR4 probes (k = 0.1 N/m). The scan rate, feedback gains, and deflection setpoint were adjusted during acquisition to optimize image quality. HS-AFM imaging was carried out in amplitude-modulation mode using a modified high-speed AFM (SS-NEX, RIBM, Tsukuba, Japan) optimized for molecular-resolution imaging[58]. The instrument was equipped with a custom digital high-speed lock-in amplifier coded on a reconfigurable FPGA platform using LabVIEW (National Instruments, Austin, TX, USA). Short cantilevers (∼7 µm) designed for HS-AFM and fitted with electron-beam-deposited (EBD) tips were used (USC-F1.2-k0.15, Nanoworld, Switzerland). These cantilevers have a nominal spring constant of 0.15 N/m, a resonance frequency in liquid of ∼500 kHz, and a quality factor (Q) of ∼2. The deflection sensitivity was 0.1 V/nm, and the imaging amplitude setpoint was ∼90% of the free amplitude (∼10 Å). All HS-AFM experiments were performed at 35 °C inside a temperature-controlled enclosure.

### Image analysis

Acquired images were plane-fit and flattened using the AFM processing and analysis software provided by the instrument manufacturer. HS AFM image treatment was limited to the correction of XY drift and a first-order X-line fit. Image analysis was performed using the general distribution of ImageJ[59] and of WSxM[60].

### Building of model assemblies

To improve accuracy and resolve ambiguities in protein position assignment from AFM images, three independent coordinate extractions were performed from the same micrographs and superimposed. Coordinates found in only one of the three datasets were discarded, while the remaining positions were averaged. This procedure yielded an overall precision of 5.7 ± 3.8 Å for the picked protein centers. These coordinates were used to place proteins and lipids extracted from a previously conducted coarse-grained MD simulation of VDAC embedded in a lipid membrane[17]. Distinct configurations were randomly selected, and proteins were extracted with surrounding cylindrical lipid shells (radius: 5 nm) centered on each protein. Overlapping lipids were sequentially removed to avoid steric clashes. The resulting structural models were analyzed to evaluate lipid-protein and inter-protein contacts, providing molecular-level insights into the assemblies. The Python program is freely available at https://github.com/CodeCanSolveAll/Particle_Analysis.

### Protein distance distribution

To quantitatively assess whether VBIT-4 affects VDAC1 organization, we analyzed protein compaction within clusters by measuring inter-protein distance distributions derived from AFM-extracted protein coordinates. The program is available at https://github.com/deejy/ComBiol2025. Nearest-neighbour relationships were identified using Delaunay triangulation, and pairwise distances were classified into discrete ranges corresponding to increasing numbers of intervening lipid layers. To compare the distributions obtained from the eight analyzed AFM fields, we used a linear mixed-effects model, with VBIT-4 concentration as a fixed effect and the AFM field (i.e., each independent set of detected protein coordinates) included as a random intercept. This approach accounts for the non-independence of multiple distance measurements derived from the same AFM field. Treating these as independent would lead to pseudo-replication and inflated statistical power. The model estimates the average inter-protein distance as a function of VBIT-4 concentration, and the reported coefficients represent the change in mean distance relative to the reference condition (no VBIT-4). Associated p-values assess whether the observed differences are large relative to the variability between fields, and therefore whether they are consistent with a genuine effect of VBIT-4 rather than random variation.

### Preparation of nanodiscs

MSP1D1 membrane scaffold protein was produced and purified as described by Ritchie, T. K. et al.[61]. Chloroform-solubilised lipids (Chicken egg phosphatidylcholine) were dried under nitrogen gas and re-solubilised in 20 mM Tris pH 8.0 buffer containing 50mM sodium cholate at a final concentration of 25 mM lipid. Lipids and MSP were mixed at a 65:1 molar ratio at 4°C for 1 hour. Bio-Beads SM2 were added at a 50x excess (weight) of total detergent, and the mix was incubated overnight at 4 °C on agitation. The nanodiscs were filtered through a 0.2 µm sterile syringe filter and centrifuged at 100,000 x g for 30 minutes at 4 °C. The supernatant was subjected to size exclusion chromatography on a Superdex 200 increase 10/300 GL column in 20 mM HEPES pH 7.4, 100 mM NaCl. The nanodisc-containing fractions were concentrated for labelling. Nanodiscs were labelled using the Protein Labelling Kit RED-NHS 2^nd^ Generation from NanoTemper (NT-L021) at a 1:1.5 protein:dye molar ratio in 20 mM HEPES buffer pH 7.4, 100 mM NaCl for 2 hours at 4°C in the dark. Excess dye was removed using a Micro Bio-Spin P6 column equilibrated in the same buffer. The degree of labelling was determined using UV-Vis spectrophotometry at 650 nm and 280 nm according to the manufacturer’s instructions. The degree of labelling was between 60-75% (ε_nanodisc =_ 36,900 M^-1^ cm^-1^, ε_dye =_ 195,000 M^-1^ cm^-1^) across the replicates.

### Preparation of VDAC1 nanodiscs

mVDAC1 was purified and refolded as explained above, except the final gel filtration was performed in 20mM HEPES buffer pH 7.4, 100mM NaCl, 0.1% LDAO. Detergent-refolded VDAC1 was labelled using the Protein Labelling Kit RED-NHS 2nd Generation from NanoTemper (NT-L021) at a 1:1.5 protein:dye molar ratio for 2 hours at 4°C in the dark. Excess dye was removed using a Micro Bio-Spin P6 column equilibrated in the gel filtration buffer. The degree of labelling was determined using UV-Vis spectrophotometry at 650 nm and 280 nm according to the manufacturer’s instructions. The degree of labelling was 30% (ε_VDAC1 =_ 43,890 M-1 cm-1, εdye = 195,000 M-1 cm-1). Labelled VDAC1 was reconstituted into MSP1D1 nanodiscs by incubating proteins with cholate-solubilised lipids (EggPC) at a 65:1 molar ratio lipids:MSP and 12:1 molar ratio MSP:VDAC1 for 1 hour at 4°C before adding 50x excess (wt) of BioBeads SM2 and incubating overnight at 4 °C on agitation. The nanodiscs were filtered through a 0.2 µm sterile syringe filter and centrifuged at 100,000 x g for 30 minutes at 4 °C. The supernatant was subjected to size exclusion chromatography on a Superdex 200 increase 10/300 GL column in 20 mM HEPES pH 7.4, 100 mM NaCl. Reconstituted VDAC1 was concentration in an Amicon-50 and used in MST experiments.

### Microscale Thermophoresis (MST)

MST experiments were performed using a NanoTemper Monolith X MO-G039, Munich, Germany. VBIT-4 (Selleckchem #S3544) was diluted in DMSO at a final concentration of 10 mM and sub-stocks of 1 mM were made in buffer 20 mM HEPES pH 7.4, 100 mM NaCl containing 10 % DMSO. 15 µL of 50nM labelled nanodiscs or VDAC1-containing nanodiscs were incubated with 15 µL of VBIT-4 (0.25 - 500 µM) and incubated for 20 minutes at room temperature as performed by Ben-Hail et. al.[17]. Following a short tabletop spin, 10 µL of the samples were loaded into 12 glass capillaries (NanoTemper premium capillaries, MO-K022) and thermophoresis was performed (60% excitation power at 650nm, and a medium infra-red laser power with a wavelength of 1480nm). The final concentration of DMSO in the capillaries was maintained at 5%. Thermophoresis experiments were performed as described by the manufacturer and data were analysed at 650 nm at a laser ON-time of 2.5s since they had the best SNR (Signal to Noise Ratio. Data were fitted by the system.

### Laurdan generalised polarisation

Laurdan-containing liposomes were prepared by mixing chloroform-solubilized lipids (POPC:POPE:Chol, 62.5:30:7.5) at a concentration of 1 mM total lipids with chloroform-solubilized laurdan at a lipid:laurdan molar ratio of 1000:1. Lipid mixtures were dried under nitrogen gas in the dark. Liposomes were formed by resuspending dried lipids in 200 µL of 20 mM HEPES buffer pH 7.4 containing 100 mM NaCl and vortexing. Liposomes were sonicated with 10 pulses at 10% power, 10% ON/OFF cycle to form SUVs with an average diameter of 100 nm as verified by dynamic light scattering. Laurdan-containing liposomes were diluted 10x in buffer and added to a quartz cuvette (pathlength 3 mm) at a final volume of 150 µL. Sub-stocks of VBIT-4 in DMSO were made such that the final volume added to the cuvette would be a constant 1 µL. Control samples contained the same volume of DMSO. The excitation wavelength was set to 375 nm and the emission scan range was from 400-600 nm. Excitation slit width – 10 nm, emission slit width – 5 nm. Data were recorded at a scanning speed of 600 nm/min at 26 °C on a Cary Eclipse fluorescence spectrometer. Generalised polarisation was calculated as GP = (I_440 –_ I_490)_ / (I_440 +_ I_490)_. GP was plotted as a function of VBIT-4 concentration on Python using a partitioning model.

### Planar lipid bilayer measurements

Planar lipid membranes (PLM) were formed by two different techniques: 1) using “dry” membranes formed by the lipid monolayer opposition technique on a circular aperture in a Teflon partition dividing two ∼1.5 mL (*cis* and *trans*) compartments of the experimental chamber, as previously described[62], and 2) using “painted” membranes from lipid solution in decane using Orbit mini system (Nanion Technologies, Minich, Germany)[63]. Lipids dioleoyl-phosphatidylcholine (DOPC), dioleoyl-phosphatidylethanolamine (DOPE), diphytanoyl-phosphatidylcholine (DPhPC), polar lipid extract from soybean (PLE), and cholesterol (Avanti Polar Lipids) were used for PLM preparations. Aqueous solutions consisted of 150 mM KCl buffered with 5 mM HEPES at pH 7.4. In the first method of PLM formation, the 5 mg/mL lipid mixtures in pentane were used. The aperture in the Teflon partition was pretreated by 1% hexadecane in pentane before PLM formation, thus leaving only a residual amount of hexadecane in the bilayers. The current recordings were performed as described previously[62], using Axopatch 200B amplifier (Axon Instruments) in the voltage-clamp mode. Data were filtered by a low-pass 8-pole Butterworth filter (Model 900 Frequency Active Filter, Frequency Devices) at 15 kHz, digitized with a sampling frequency of 50 kHz, and analysed using pClamp 10.7 software (Axon Instrument). For data analysis, digital filtering using a 1 or 0.5 kHz low-pass Bessel filter was applied. Potential is defined as positive when it is greater on the *cis* side, the side of VBIT-4 and VDAC1 addition. In the second method, lipid bilayers were formed the Orbit mini system (Nanion Technologies, Munich, Germany) by painting the chips (Ionera, Freiburg, Germany) with the 5 mg/mL lipid mixtures in decane. Multi Electrode Cavity Array (MECA4) chips were filled with 150 μL of 150 mM KCl solution at pH 7.4. VBIT-4 in DMSO solution was added to the bilayer in both experimental setups after PLM was formed, up to a total volume of less than 5 μL, and PLM conductance of less than 0.1 pS was verified. Addition of the corresponding DMSO aliquots to the membrane-bathing solutions did not change membrane conductance[64]. All measurements were performed at room temperature of 21 ± 1°C.

### VDAC conductance measurements

VDAC1 channel insertion was achieved by adding 0.1-1.5 μL of VDAC1 diluted in buffer containing 10 mM Tris, 50 mM KCl, 1 mM EDTA, 15 % (v/v) DMSO, 2.5% (v/v) Triton X-100, pH 7.0. Current recording and its analysis were performed as described previously[65]. VDAC1 voltage-gating properties were assessed following the previously described protocol [66] under the application of a slow symmetrical 5 mHz triangular voltage wave of ±60 mV amplitude from an Arbitrary Waveform Generator 33220A (Agilent). Data were acquired at a sampling frequency of 2 Hz and analyzed as described previously[66] using an algorithm developed in-house and pClamp 10.7 software. In each experiment, current records were collected from membranes containing 10-50 channels in response to 5–10 periods of voltage waves to ensure data collection from more than 50 channels per experiment. Only the part of the wave, during which the channels were reopening, was used for the subsequent analysis[67].

### Liposome leakage assay

Liposomes (POPC:POPE:chol, 62.5:30:7.5 w:w) were prepared by mixing 5mg of chloroform-solubilized lipids in a glass tube. Dried lipids were resuspended in 200 µL of 100 mM fluorescein dissolved in HEPES buffer pH 7.4, and vortexed vigorously. Liposomes were subjected to 3 cycles of freeze-thawing in liquid nitrogen followed by 50°C in a Thermomix. Liposomes were then sonicated with 20 pulses at 20% power, 10% ON/OFF cycle to form SUVs of diameter 50-100 nm as verified by dynamic light scattering. Liposomes were pelleted by centrifugation at 25,000 rpm for 30 mins at 4°C on a TLA55 rotor. The supernatant was discarded, and liposomes were washed with 1 mL of HEPES buffer. The washing/centrifugation step was repeated till no free fluorescein was detectible by absorbance spectroscopy. Pelleted liposomes were resuspended in 1 mL of HEPES buffer, and 32 µL of liposomes were used in each 200 µL reaction in a 96-well plate, giving a final concentration of 1 mM lipid/reaction. Sub-stocks of VBIT-4 in DMSO were made such that the final volume added to each well would be a constant 1 µL. The negative control sample included 1 µL of DMSO, while the positive control sample included 1 µL of 100% Triton X-100. Time-resolved fluorescence spectra were recorded in a Tecan Spark plate reader for 2 hours every 30 seconds with shaking between each reading. Excitation and emission wavelengths for fluorescein were set to 485 nm and 535 nm, respectively. At the end of the kinetics, 1 μL of Triton X-100 was added to solubilise the liposomes. Data were normalised w.r.t. liposomes treated with DMSO as F_norm =_ (F-F_min)_/(F_max –_ F_min)_ after 100 min, and a histogram of % release was plotted.

### VBIT-4 CG parameterization

### VBIT-4 charge state

VBIT-4 contains a piperazine group with two nitrogen atoms. These are typically protonable under physiological conditions, with pKa values usually in the range of 7–9. Out of these two, the most protonable nitrogen is the one closer to the hydroxyl (-OH) and amide (-CONH-) groups. The nitrogen near the benzene ring is both sterically hindered and electronically less basic due to the -OCF_3 g_roup’s electron-withdrawing effect. The nitrogen near the amide and hydroxyl is both more exposed and more basic, making it the preferred protonation site. Given the significant conformational differences observed between the neutral and protonated versions of VBIT-4 in our reference AA simulations, we developed a Martini CG model for each state and analyzed their effects separately.

### All-atom reference simulations

VBIT-4 Martini 3 models were based on simulations of all-atom counterparts, parameterized using the OPLS-AA force field via LigParGen[68–70]. Each system was simulated for 500 ns in water. Each system was minimized using steepest descent with 5000 steps, followed by a relaxation of 250 ps using a 1 fs time step, and a production run of 500 ns using a 2 fs time step. The temperature and pressure were held constant at 298 K and 1 bar using the Berendsen thermostat and barostat[71] for the relaxation step and the V-rescale thermostat[72] and Parrinello-Rahman barostat[73] for the production run. The pressure coupling was isotropic with a compressibility of 4.5 × 10^−5^ bar^−1^. The temperature was coupled using *τ* = 1 ps during both the relaxation and production run. For pressure coupling, *τ* = 1 ps was set during relaxation and 5 ps during production. The electrostatics were treated with the PME algorithm[74] using a cutoff of 1.2 nm, while the van der Waals interactions were truncated, smoothly switching the force to zero between 1.0 and 1.2 nm[75]. Bonds involving hydrogen atoms were restricted using the LINCS algorithm[76].

### Coarse-grained model parameterization

Martini CG models were then obtained according to the parametrization rules of Martini 3, as described elsewhere[77]. In short, atomistic simulations were mapped into CG resolution using MD Analysis[78, 79] . The bonded terms were then measured and the distributions plotted. The molecule was then modelled in CG and simulated in water. Bonded distributions were fitted onto the corresponding atomistic distributions. Bead types and sizes were selected according to Martini 3 parametrization guidelines, ensuring consistency with default bead assignments for specific chemical groups[77]. *These choices were validated by calculating CG octanol/water partitioning free energies for neutral VBIT and comparing them against a consensus of reference values obtained by theoretical predictors. Rather than comparing our Martini logP measurements against atomistic molecular dynamics calculations, we benchmark against established logP prediction methods. For equilibrium octanol/water partitioning, empirical predictors have generally demonstrated accuracy comparable to, or better than, atomistic free-energy calculations when evaluated against experiment[80]. We further use a consensus of five empirical models, as consensus predictions have been shown to outperform individual predictors[81]. Reference logP values for neutral VBIT were obtained using the prediction methods available through SwissADME[82] (iLOGP[83], XLOGP3[84], WLOGP[85], MLOGP[86], SILICOS-IT), yielding predicted logP values of 3.66, 3.85, 4.06, 2.25, and 3.28, respectively. The average of these predictions was used as a consensus estimate, giving a logP value of 3.42 ± 0.63*.

*The calculated CG partitioning free energies were obtained for neutral VBIT by thermodynamic integration as described elsewhere[77]. In short, the solute is alchemically decoupled from each solvent environment independently across 12 lambda windows, gradually turning off all non-bonded interactions between the solute and its surroundings. The free energy change along this path corresponds to the solvation free energy in that solvent, and taking the difference between octanol and water (ΔG_octanol − ΔG_water) yields the transfer free energy, which is converted to a partition coefficient. Three independent replicates were run, yielding octanol-water logP values of 3.54, 3.53, and 3.51, with an average of 3.52 ± 0.01. Convergence and overlap analysis confirmed that all lambda windows were well-sampled and that the free energy estimates were statistically reliable as required per the guidelines for the analysis of free energy calculations[87]; the corresponding forward/backward convergence plots and MBAR overlap matrices are provided in the Supplementary figures 11 and 12*.

### VBIT-4 Martini CG MD simulations

*Unbiased CG MD.* Membrane bilayers were constructed with COBY[88], which also solvated the system, neutralized it, and added ions to obtain a 150 mM NaCl concentration. COBY also added the appropriate number of VBIT-4 molecules in solution. All systems were subject to the same relaxation and production protocol. First, a minimization of 5000 steps using steepest descent, followed by 10 ns of relaxation, and finally at least 5 *μ*s production run. A complex membrane composition mimicking the mitochondrial outer membrane was used

(8:25:17:13:4:9:3:8:4 CHOL:POPC:DOPC:DPPC:POPE:DOPE:DPPE:POPS:POPI:DPPI) in a 15 × 15 × 17.5 nm^3^ system. Prior to production, the systems underwent 5000 energy minimization steps, followed by equilibration for 10 ns using a 10 fs time step. During equilibration, VBIT-4 molecules were pulled into the membrane using flat-bottom potentials to avoid sampling the membrane adsorption process. These potentials were not applied during the production run. The production runs for each system were carried out for 10 µs. Throughout the simulations, the temperatures of the lipids (including VBIT-4) and solvent (including ions) were independently maintained at 300 K using the v-rescale thermostat (τ_t =_ 1 ps)[72], and the pressure was kept at 1 bar using the c-rescale barostat (τ_p =_ 4 ps)[89] with semi-isotropic pressure coupling applied during both equilibration and production phases. Compressibility was set to 3 × 10^-4^ bar^-1^. As suggested by Kim et al.[90], the cut-off distance for the short-range neighbor list was set to 1.35 nm for proper neighbor list updates.

### Membrane translocation PMF

Similarly to the unbiased CG MD systems, COBY was used to build a smaller 7 × 7 × 13.5 nm^3^ system, fully solvated with water and 150 mM NaCl. One VBIT-4 molecule was placed 4 nm away (determined from the center of geometry of VBIT-4) from the bilayer center along the bilayer normal, and overlapping water molecules were removed. Minimization and equilibration were performed as with the unbiased simulations. At the end of the equilibration, initial configurations for the umbrella sampling windows were generated by pulling the VBIT-4 molecule through the bilayer center at a rate of 0.0001 nm/ps and a force constant of 3000 kJ/mol/nm^2^. In each umbrella sampling window, a harmonic potential was applied to keep the VBIT-4 molecule in its pre-defined distance from the bilayer center with a force constant ranging from 1000 to 2000 kJ/mol/nm^2^. A total of 52 windows were simulated for 2 μs and the data from the first 200 ns were discarded before applying the weighted histogram analysis method to estimate the free energy profile using GROMACS. The translocation PMF of the molecule was computed along the entire membrane normal from z = 5.25 to z = -5.25. For each replica, the PMF was symmetrized by splitting it at the membrane center and averaging both halves. The final PMF and standard deviation were obtained by averaging the symmetrized profiles across all replicas.

### Polar defect formation PMF

Polar defect formation PMF was determined as described by Hub and Awasthi[42]. Small 10 × 10 × 14 nm^3^ membrane bilayer systems were built using COBY, as described for the unbiased MD systems. Minimization and equilibration were performed as with the unbiased simulations. The initial frames for the umbrella sampling simulations were extracted from constant velocity pulling simulations. In these simulations, the systems were pulled from the reaction chain coordinate[42, 91] *ξ*_ch =_ 0.25 to *ξ*_ch =_ 1 over 1000 ns using a force constant of 5000 kJ mol^-1^. Nineteen umbrella windows were employed, with reference positions ranging from *ξ*ch = 0.1 to 1 in increments of 0.05, where 1 is the fully formed pore. The force constant was set to 8000 kJ mol^-1^. Each window was simulated for 1 *μ*s, with the initial 500 ns excluded for equilibration. Three replicas of each window were run. The potential of mean force was computed using the weighted histogram analysis method (WHAM)[92].

### VDAC oligomerisation CG MD simulations

System construction followed the same protocol as the unbiased CG MD simulations described above. Briefly, COBY was used to build a 30 × 30 × 13.5 nm³ system containing a lipid bilayer with a composition mimicking the mitochondrial outer membrane, fully solvated with water and 150 mM NaCl. Nine VDAC-1 copies were inserted into the membrane in an equidistant 3 × 3 grid. The VDAC-1 CG model was built using martinize2[93] from a reference NMR structure (PDB: 6TIQ)[94] with default settings and an elastic network tertiary bias (force constant: 700 kJ/mol; cutoff: 0.85 nm). Where applicable, VBIT-4 molecules were added in solution by COBY, with an exclusion radius of 3 Å around each VDAC-1 protein to prevent placement in the immediate protein neighbourhood. All systems followed the same relaxation and production protocol as the unbiased CG MD simulations, with the addition of a third temperature coupling group for the VDAC-1 proteins. Like before, during equilibration, VBIT-4 molecules were pulled into the membrane using flat-bottom potentials to avoid sampling the membrane adsorption process. Ten replicas were prepared per system with identical starting structures but different random seeds, each simulated for 10 μs.

### Oligomerisation analysis

VDAC oligomerisation was quantified from simulation trajectories using a DBSCAN clustering approach[95] implemented in sklearn[96]. For each trajectory frame, the centre of mass of each VDAC protein was computed and a pairwise distance matrix was constructed accounting for periodic boundary conditions. DBSCAN (min_samples=1, ε=55 Å) was applied to assign proteins to clusters. The ε cutoff was derived from the centre-of-mass distance distribution of a fully oligomerised VDAC-only trajectory. The fraction of proteins belonging to the largest cluster was used as the primary observable, providing a continuous measure of oligomerisation extent ranging from 1/N (all proteins isolated) to 1.0 (all proteins in a single assembly), where N is the total number of VDAC copies.

Two complementary metrics were derived per replicate. The observed oligomerisation extent was taken as the fraction in the largest cluster at the final simulation frame, reflecting the degree of assembly reached within the simulated timescale. The predicted oligomerisation plateau was estimated by fitting a hyperbolic model f(t) = L·t / (t₅₀ + t) to each trajectory, where L is the asymptotic plateau and t₅₀ is the half-time. Prior to fitting, trajectories were smoothed using a uniform filter (window = 5% of trajectory length) and data points were weighted proportionally to their smoothed value, down-weighting early stochastic association events and emphasising the plateau region. The fitted plateau L was constrained to [f₀, 1.0], where f₀ is the initial value.

All pairwise comparisons between conditions were performed using the two-sided Mann-Whitney U test, chosen for its robustness to non-normality given the small sample size (n = 10 replicates per condition). P-values were corrected for multiple comparisons using the Holm stepwise procedure across the three pairwise comparisons per metric. Effect sizes are reported as rank-biserial correlations. Significance thresholds: \*p < 0.05, \*\*p < 0.01, \*\*\*p < 0.001, \*\*\*\*p < 0.0001.

### Cell toxicity assay

HeLa WT and VDAC1 knockout cells from Synthego were seeded at 2,500 cells/well in a 96-well plate (ThermoFisher Scientific, 165305) in DMEM (Gibco, 15607) supplemented with 10% FBS (Gibco, 10437). VBIT-4 was diluted in the above media (pH 7.4) and added to the cells at a final concentration of 0-200 µM VBIT-4. After 48 hours at 37 °C and 5% CO_2,_ GF-AFC and bis-AAF-R110 substrates from the MultiTox-Fluor Multiplex Cytotoxicity Assay kit (Promega, G9201) were added to the cells following the manufacturer’s protocol and incubated for 30 minutes at 37 °C. Viability and cytotoxicity were measured using 400/505 nm and 485/520 nm excitation/emission filters, respectively, in a CLARIOstar plate reader. Nonlinear regression curve fit was applied to calculate the IC_50 f_or VBIT-4 using GraphPad Prism (version 11.0.0).

### Mitochondrial oxygen consumption rate measurement

HeLa cells were seeded in a 96-well plate at 10,000 cells/well in DMEM with 10% FBS and the oxygen consumption rate (OCR) were measured with Resipher device (Lucid Scientific, Atlanta, GA, USA). Baseline OCR was measured overnight before treatment with vehicle control, 1.25 to 10 µM VBIT-4, 0.5 µM oligomycin, 2 µM and 4 µM FCCP. OCR data after 6 hours of incubation with drugs were normalized to the baseline OCR for each well.

### Mitochondrial mass, membrane potential, and calcium measurements by flow cytometry

HeLa cells were seeded into 24-well plates at 1×10^5^ cells/well in DMEM with 10% FBS. Cells were mock-treated or treated with 5 µM or 10 µM VBIT-4 overnight (16 hours). Trypsinized cells were washed and loaded with Mitotracker Green for flow cytometry (MTG, 0.5 µl/ml for mitochondrial mass) and tetramethylrhodamine methyl ester (TMRM, 10 nM, for membrane potential) in PBS for 30 minutes at 37 °C with Live/Dead Near-IR dye added for the last 15 minutes. Alternatively, cells were stained with Rhod2-AM (1 µM, calcium indicator) and 0.02% Pluronic F-127 in PBS for 1 hour at room temperature, washed and resuspended in 250 µL of PBS and incubated at room temperature for 30 minutes with Live/Dead Near-IR dye added for the last 15 minutes (all reagents from Thermo Fisher). Cells were acquired on a BD FACSymphony A5 with TMRM or Rhod2 signal acquired in YG 586/15, MTG acquired in Blue 530/30, and live cell dye in Red 780/60. Analysis was performed in FlowJo 10.10 (Becton Dickinson) with cell gate established by forward scatter versus side scatter, followed by singlet gate, live cell gate, and determination of mean fluorescence intensity of MTG, TMRM, or Rhod2.

### FTIR

A Bruker TENSOR FT-IR spectrophotometer equipped with an ATR plate and an MCT D316 nitrogen-cooled detector was used to further characterize the VBIT-4 IR absorption band and the pH-dependent solubility of the molecule. A VBIT-4 suspension containing 10 µL of VBIT at 25 mg/mL in DMSO, 20 µL of H₂O, and 10 µL of HCl (0.03%) is prepared just prior to the experiment. Four µL of the slightly turbid suspension is deposited on the surface of the diamond plate and equilibrated for 15 minutes with a cover slip to avoid evaporation. At this point, the upper liquid is adsorbed using the capillarity of a cleaning paper, and the surface is washed twice using 4 µL of 50 mM phosphate buffer (pH 7.6) and 100 mM NaCl.

This procedure is sufficient to completely remove the FTIR signal of DMSO and shows a strong adsorption of VBIT-4 on the diamond surface, as depicted by two characteristic areas of absorption bands in the 1470-1570 cm⁻¹ and 1125-1300 cm⁻¹ regions. From this starting situation, we characterize the tendency of the adsorbed VBIT-4 to elute using phosphate buffer (pH 7.6).

## Acknowledgements

We thank Dr. Manuel N. Melo for the insightful suggestion that inspired the polar defect formation simulations. This work was supported by the Aix-Marseille Université grant AMX-21-PEP-022 (L.B.) and a PhD fellowship from Aix-Marseille Université (V.R.). L.B.A. acknowledges the support of the French National Centre for Scientific Research and the HPC resources provided by IDRIS, CINES, and TGCC through GENCI (allocation 2024-A0160713456). L.B.A. also acknowledges the support of the Centre Blaise Pascal de Simulation et Modélisation Numérique’s IT test platform at ENS de Lyon (Lyon, France) for the computer facilities. M.S.W., W.F., N.A.B., B.G.B., M.G.L., M.R., S.M.B., and T.K.R. acknowledge the support of the Intramural Research Program of the National Institutes of Health (NIH), *Eunice Kennedy Shriver* National Institute of Child Health and Human Development (NICHD). This research was supported in part by the Intramural Research Program of the National Institutes of Health (NIH). The contributions of the NIH authors are considered Works of the United States Government. The findings and conclusions presented in this paper are those of the authors and do not necessarily reflect the views of the NIH or the U.S. Department of Health and Human Services.

## References

1. Gonçalves RP, Buzhynskyy N, Prima V, Sturgis JN, Scheuring S (2007) Supramolecular assembly of VDAC in native mitochondrial outer membranes. J Mol Biol 369:413–418. 10.1016/j.jmb.2007.03.063

2. Hoogenboom BW, Suda K, Engel A, Fotiadis D (2007) The supramolecular assemblies of voltage-dependent anion channels in the native membrane. J Mol Biol 370:246–255. 10.1016/j.jmb.2007.04.073

3. Naghdi S, Várnai P, Hajnóczky G (2015) Motifs of VDAC2 required for mitochondrial Bak import and tBid-induced apoptosis. Proc Natl Acad Sci U S A 112:E5590–5599. 10.1073/pnas.1510574112

4. Yuan Z, Dewson G, Czabotar PE, Birkinshaw RW (2021) VDAC2 and the BCL-2 family of proteins. Biochem Soc Trans 49:2787–2795. 10.1042/BST20210753

5. Sander P, Gudermann T, Schredelseker J (2021) A Calcium Guard in the Outer Membrane: Is VDAC a Regulated Gatekeeper of Mitochondrial Calcium Uptake? Int J Mol Sci 22:946. 10.3390/ijms22020946

6. Shankar TS, Ramadurai DKA, Steinhorst K, Sommakia S, Badolia R, Thodou Krokidi A, Calder D, Navankasattusas S, Sander P, Kwon OS, Aravamudhan A, Ling J, Dendorfer A, Xie C, Kwon O, Cheng EHY, Whitehead KJ, Gudermann T, Richardson RS, Sachse FB, Schredelseker J, Spitzer KW, Chaudhuri D, Drakos SG (2021) Cardiac-specific deletion of voltage dependent anion channel 2 leads to dilated cardiomyopathy by altering calcium homeostasis. Nat Commun 12:4583. 10.1038/s41467-021-24869-0

7. Rosencrans WM, Rajendran M, Bezrukov SM, Rostovtseva TK (2021) VDAC regulation of mitochondrial calcium flux: From channel biophysics to disease. Cell Calcium 94:102356. 10.1016/j.ceca.2021.102356

8. Queralt-Martín M, Bergdoll L, Teijido O, Munshi N, Jacobs D, Kuszak AJ, Protchenko O, Reina S, Magrì A, De Pinto V, Bezrukov SM, Abramson J, Rostovtseva TK (2020) A lower affinity to cytosolic proteins reveals VDAC3 isoform-specific role in mitochondrial biology. J Gen Physiol 152:. 10.1085/jgp.201912501

9. Xu X, Decker W, Sampson MJ, Craigen WJ, Colombini M (1999) Mouse VDAC isoforms expressed in yeast: channel properties and their roles in mitochondrial outer membrane permeability. J Membr Biol 170:89–102. 10.1007/s002329900540

10. Camara AKS, Zhou Y, Wen P-C, Tajkhorshid E, Kwok W-M (2017) Mitochondrial VDAC1: A Key Gatekeeper as Potential Therapeutic Target. Front Physiol 8:460. 10.3389/fphys.2017.00460

11. Magri A, Messina A (2017) Interactions of VDAC with Proteins Involved in Neurodegenerative Aggregation: An Opportunity for Advancement on Therapeutic Molecules. Curr Med Chem 24:4470–4487. 10.2174/0929867324666170601073920

12. Reina S, De Pinto V (2017) Anti-Cancer Compounds Targeted to VDAC: Potential and Perspectives. Curr Med Chem 24:4447–4469. 10.2174/0929867324666170530074039

13. Kim J, Gupta R, Blanco LP, Yang S, Shteinfer-Kuzmine A, Wang K, Zhu J, Yoon HE, Wang X, Kerkhofs M, Kang H, Brown AL, Park S-J, Xu X, Zandee van Rilland E, Kim MK, Cohen JI, Kaplan MJ, Shoshan-Barmatz V, Chung JH (2019) VDAC oligomers form mitochondrial pores to release mtDNA fragments and promote lupus-like disease. Science 366:1531–1536. 10.1126/science.aav4011

14. Jahn H, Bartoš L, Dearden GI, Dittman JS, Holthuis JCM, Vácha R, Menon AK (2023) Phospholipids are imported into mitochondria by VDAC, a dimeric beta barrel scramblase. Nat Commun 14:8115. 10.1038/s41467-023-43570-y

15. Leung MR, Zenezini Chiozzi R, Roelofs MC, Hevler JF, Ravi RT, Maitan P, Zhang M, Henning H, Bromfield EG, Howes SC, Gadella BM, Heck AJR, Zeev-Ben-Mordehai T (2021) In-cell structures of conserved supramolecular protein arrays at the mitochondria-cytoskeleton interface in mammalian sperm. Proc Natl Acad Sci U S A 118:e2110996118. 10.1073/pnas.2110996118

16. Callegari S, Kirk NS, Gan ZY, Dite T, Cobbold SA, Leis A, Dagley LF, Glukhova A, Komander D (2025) Structure of human PINK1 at a mitochondrial TOM-VDAC array. Science 388:303–310. 10.1126/science.adu6445

17. Lafargue E, Duneau J-P, Buzhinsky N, Ornelas P, Ortega A, Ravishankar V, Sturgis J, Casuso I, Bergdoll L (2025) Membrane lipid composition modulates the organization of VDAC1, a mitochondrial gatekeeper. Commun Biol 8:936. 10.1038/s42003-025-08311-5

18. Mannella CA, Colombini M, Frank J (1983) Structural and functional evidence for multiple channel complexes in the outer membrane of Neurospora crassa mitochondria. Proc Natl Acad Sci U S A 80:2243–2247. 10.1073/pnas.80.8.2243

19. Ben-Hail D, Begas-Shvartz R, Shalev M, Shteinfer-Kuzmine A, Gruzman A, Reina S, De Pinto V, Shoshan-Barmatz V (2016) Novel Compounds Targeting the Mitochondrial Protein VDAC1 Inhibit Apoptosis and Protect against Mitochondrial Dysfunction. J Biol Chem 291:24986–25003. 10.1074/jbc.M116.744284

20. Klapper-Goldstein H, Verma A, Elyagon S, Gillis R, Murninkas M, Pittala S, Paul A, Shoshan-Barmatz V, Etzion Y (2020) VDAC1 in the diseased myocardium and the effect of VDAC1-interacting compound on atrial fibrosis induced by hyperaldosteronism. Sci Rep 10:22101. 10.1038/s41598-020-79056-w

21. Verma A, Shteinfer-Kuzmine A, Kamenetsky N, Pittala S, Paul A, Nahon Crystal E, Ouro A, Chalifa-Caspi V, Pandey SK, Monsonego A, Vardi N, Knafo S, Shoshan-Barmatz V (2022) Targeting the overexpressed mitochondrial protein VDAC1 in a mouse model of Alzheimer’s disease protects against mitochondrial dysfunction and mitigates brain pathology. Transl Neurodegener 11:58. 10.1186/s40035-022-00329-7

22. Belosludtsev KN, Serov DA, Ilzorkina AI, Starinets VS, Dubinin MV, Talanov EY, Karagyaur MN, Primak AL, Belosludtseva NV (2023) Pharmacological and Genetic Suppression of VDAC1 Alleviates the Development of Mitochondrial Dysfunction in Endothelial and Fibroblast Cell Cultures upon Hyperglycemic Conditions. Antioxidants (Basel) 12:1459. 10.3390/antiox12071459

23. Mohammad Al-Amily I, Sjögren M, Duner P, Tariq M, Wollheim CB, Salehi A (2023) Ablation of GPR56 Causes β-Cell Dysfunction by ATP Loss through Mistargeting of Mitochondrial VDAC1 to the Plasma Membrane. Biomolecules 13:557. 10.3390/biom13030557

24. Wei S-N, Zhang H, Lu Y, Yu H-J, Ma T, Wang S-N, Yang K, Tian M-L, Huang A-H, Wang W, Li F-S, Li Y-W (2023) Microglial voltage-dependent anion channel 1 signaling modulates sleep deprivation-induced transition to chronic postsurgical pain. Sleep 46:zsad039. 10.1093/sleep/zsad039

25. Luo X, Yue J (2025) VDAC1 Inhibition Mitigates Inflammatory Status and Oxidative Stress in Epileptic Mice Treated with the Ketogenic Diet. Neurochem Res 50:118. 10.1007/s11064-025-04366-2

26. Terrones O, Antonsson B, Yamaguchi H, Wang H-G, Liu J, Lee RM, Herrmann A, Basañez G (2004) Lipidic Pore Formation by the Concerted Action of Proapoptotic BAX and tBID. Journal of Biological Chemistry 279:30081–30091. 10.1074/jbc.M313420200

27. Colombini M (2019) Ceramide Channels. In: Stiban J (ed) Bioactive Ceramides in Health and Disease. Springer International Publishing, Cham, pp 33–48

28. Anosov AA, Smirnova EYu, Ryleeva ED, Gligonov IA, Korepanova EA, Sharakshane AA (2020) Estimation of the parameters of the Smoluchowski equation describing the occurrence of pores in a bilayer lipid membrane under soft poration. Eur Phys J E 43:66. 10.1140/epje/i2020-11989-0

29. Steringer JP, Bleicken S, Andreas H, Zacherl S, Laussmann M, Temmerman K, Contreras FX, Bharat TAM, Lechner J, Müller H-M, Briggs JAG, García-Sáez AJ, Nickel W (2012) Phosphatidylinositol 4,5-Bisphosphate (PI(4,5)P2)-dependent Oligomerization of Fibroblast Growth Factor 2 (FGF2) Triggers the Formation of a Lipidic Membrane Pore Implicated in Unconventional Secretion. Journal of Biological Chemistry 287:27659–27669. 10.1074/jbc.M112.381939

30. Ashrafuzzaman M, Tseng C, Duszyk M, Tuszynski JA (2012) Chemotherapy Drugs Form Ion Pores in Membranes Due to Physical Interactions with Lipids. Chem Biol Drug Des 80:992–1002. 10.1111/cbdd.12060

31. Perini DA, Aguilella-Arzo M, Alcaraz A, Perálvarez-Marín A, Queralt-Martín M (2022) Dynorphin A induces membrane permeabilization by formation of proteolipidic pores. Insights from electrophysiology and computational simulations. Computational and Structural Biotechnology Journal 20:230–240. 10.1016/j.csbj.2021.12.021

32. Serra-Batiste M, Ninot-Pedrosa M, Bayoumi M, Gairí M, Maglia G, Carulla N (2016) Aβ42 assembles into specific β-barrel pore-forming oligomers in membrane-mimicking environments. Proc Natl Acad Sci USA 113:10866–10871. 10.1073/pnas.1605104113

33. Souza PCT, Alessandri R, Barnoud J, Thallmair S, Faustino I, Grünewald F, Patmanidis I, Abdizadeh H, Bruininks BMH, Wassenaar TA, Kroon PC, Melcr J, Nieto V, Corradi V, Khan HM, Domański J, Javanainen M, Martinez-Seara H, Reuter N, Best RB, Vattulainen I, Monticelli L, Periole X, Tieleman DP, de Vries AH, Marrink SJ (2021) Martini 3: a general purpose force field for coarse-grained molecular dynamics. Nat Methods 18:382–388. 10.1038/s41592-021-01098-3

34. Bieker S, Timme M, Woge N, Hassan DG, Brown CM, Marrink SJ, Melo MN, Holthuis JCM (2024) Hexokinase-I directly binds to a charged membrane-buried glutamate of mitochondrial VDAC1 and VDAC2. 2024.07.22.604557

35. Dadsena S, Bockelmann S, Mina JGM, Hassan DG, Korneev S, Razzera G, Jahn H, Niekamp P, Müller D, Schneider M, Tafesse FG, Marrink SJ, Melo MN, Holthuis JCM (2019) Ceramides bind VDAC2 to trigger mitochondrial apoptosis. Nat Commun 10:1832. 10.1038/s41467-019-09654-4

36. Mazzobre MF, Román MV, Mourelle AF, Corti HR (2005) Octanol-water partition coefficient of glucose, sucrose, and trehalose. Carbohydr Res 340:1207–1211. 10.1016/j.carres.2004.12.038

37. Fornasier F, Souza LMP, Souza FR, Reynaud F, Pimentel AS (2020) Lipophilicity of Coarse-Grained Cholesterol Models. J Chem Inf Model 60:569–577. 10.1021/acs.jcim.9b00830

38. Avdeef A (1993) pH-metric log P. II: Refinement of partition coefficients and ionization constants of multiprotic substances. J Pharm Sci 82:183–190. 10.1002/jps.2600820214

39. MolGpka: A Web Server for Small Molecule pKa Prediction Using a Graph-Convolutional Neural Network | Journal of Chemical Information and Modeling. https://pubs.acs.org/doi/abs/10.1021/acs.jcim.1c00075. Accessed 3 Dec 2025

40. Creyf HS, Van Poucke LC (1972) The free energy, enthalpy and entropy changes of the dissociation of diprotonated diamines. Thermochimica Acta 4:485–491. 10.1016/0040-6031(72)85039-1

41. Pedersen KB, Ingólfsson HI, Ramirez-Echemendia DP, Borges-Araújo L, Andreasen MD, Empereur-Mot C, Melcr J, Ozturk TN, Bennett DW, Kjølbye LR, others (2024) The Martini 3 lipidome: expanded and refined parameters improve lipid phase behavior

42. Hub JS, Awasthi N (2017) Probing a Continuous Polar Defect: A Reaction Coordinate for Pore Formation in Lipid Membranes. J Chem Theory Comput 13:2352–2366. 10.1021/acs.jctc.7b00106

43. Belosludtsev KN, Serov DA, Ilzorkina AI, Starinets VS, Dubinin MV, Talanov EY, Karagyaur MN, Primak AL, Belosludtseva NV (2023) Pharmacological and Genetic Suppression of VDAC1 Alleviates the Development of Mitochondrial Dysfunction in Endothelial and Fibroblast Cell Cultures upon Hyperglycemic Conditions. Antioxidants (Basel) 12:1459. 10.3390/antiox12071459

44. Belosludtsev KN, Ilzorkina AI, Matveeva LA, Chulkov AV, Semenova AA, Dubinin MV, Belosludtseva NV (2024) Effect of VBIT-4 on the functional activity of isolated mitochondria and cell viability. Biochim Biophys Acta Biomembr 1866:184329. 10.1016/j.bbamem.2024.184329

45. Taniguchi H, Chakraborty S, Takahashi N, Banerjee A, Caeser R, Zhan YA, Tischfield SE, Chow A, Nguyen EM, Villalonga ÁQ, Manoj P, Shah NS, Rosario S, Hayatt O, Qu R, de Stanchina E, Chan J, Mukae H, Thomas A, Rudin CM, Sen T (2024) ATR inhibition activates cancer cell cGAS/STING-interferon signaling and promotes antitumor immunity in small-cell lung cancer. Science Advances 10:eado4618. 10.1126/sciadv.ado4618

46. Bround MJ, Havens JR, York AJ, Sargent MA, Karch J, Molkentin JD (2023) ANT-dependent MPTP underlies necrotic myofiber death in muscular dystrophy. Science Advances 9:eadi2767. 10.1126/sciadv.adi2767

47. Zhang E, Al-Amily IM, Mohammed S, Luan C, Asplund O, Ahmed M, Ye Y, Ben-Hail D, Soni A, Vishnu N, Bompada P, Marinis YD, Groop L, Shoshan-Barmatz V, Renström E, Wollheim CB, Salehi A (2019) Preserving Insulin Secretion in Diabetes by Inhibiting VDAC1 Overexpression and Surface Translocation in β Cells. Cell Metabolism 29:64–77.e6. 10.1016/j.cmet.2018.09.008

48. Gulen MF, Samson N, Keller A, Schwabenland M, Liu C, Glück S, Thacker VV, Favre L, Mangeat B, Kroese LJ, Krimpenfort P, Prinz M, Ablasser A (2023) cGAS-STING drives ageing-related inflammation and neurodegeneration. Nature 620:374–380. 10.1038/s41586-023-06373-1

49. Baik SH, Ramanujan VK, Becker C, Fett S, Underhill DM, Wolf AJ (2023) Hexokinase dissociation from mitochondria promotes oligomerization of VDAC that facilitates NLRP3 inflammasome assembly and activation. Science Immunology 8:eade7652. 10.1126/sciimmunol.ade7652

50. Keinan N, Tyomkin D, Shoshan-Barmatz V (2010) Oligomerization of the mitochondrial protein voltage-dependent anion channel is coupled to the induction of apoptosis. Mol Cell Biol 30:5698–5709. 10.1128/MCB.00165-10

51. Bergdoll LA, Lerch MT, Patrick JW, Belardo K, Altenbach C, Bisignano P, Laganowsky A, Grabe M, Hubbell WL, Abramson J (2018) Protonation state of glutamate 73 regulates the formation of a specific dimeric association of mVDAC1. Proc Natl Acad Sci USA 115:E172–E179. 10.1073/pnas.1715464115

52. Schredelseker J, Paz A, López CJ, Altenbach C, Leung CS, Drexler MK, Chen J-N, Hubbell WL, Abramson J (2014) High resolution structure and double electron-electron resonance of the zebrafish voltage-dependent anion channel 2 reveal an oligomeric population. J Biol Chem 289:12566–12577. 10.1074/jbc.M113.497438

53. Takeda H, Shinoda S, Goto C, Tsutsumi A, Sakaue H, Zhang C, Hirashima T, Konishi Y, Ono H, Yamamori Y, Tomii K, Shiino H, Tamura Y, Zuttion S, Senger B, Friant S, Becker HD, Araiso Y, Kobayashi N, Kodera N, Kikkawa M, Endo T (2025) Oligomer-based functions of mitochondrial porin. Nat Commun 16:. 10.1038/s41467-025-62021-4

54. Oflaz FE, Bondarenko AI, Trenker M, Waldeck-Weiermair M, Gottschalk B, Bernhart E, Koshenov Z, Radulović S, Rost R, Hirtl M, Pilic J, Karunanithi Nivedita A, Sagintayev A, Leitinger G, Brachvogel B, Summerauer S, Shoshan-Barmatz V, Malli R, Graier WF (2025) Annexin A5 controls VDAC1-dependent mitochondrial Ca2+ homeostasis and determines cellular susceptibility to apoptosis. EMBO J 44:3413–3447. 10.1038/s44318-025-00454-9

55. Pereira-Leite C, Lopes-de-Campos D, Fontaine P, Cuccovia IM, Nunes C, Reis S (2019) Licofelone-DPPC Interactions: Putting Membrane Lipids on the Radar of Drug Development. Molecules 24:516. 10.3390/molecules24030516

56. Lopes-de-Campos D, Pereira-Leite C, Fontaine P, Coutinho A, Prieto M, Sarmento B, Jakobtorweihen S, Nunes C, Reis S (2021) Interface-Mediated Mechanism of Action-The Root of the Cytoprotective Effect of Immediate-Release Omeprazole. J Med Chem 64:5171–5184. 10.1021/acs.jmedchem.1c00251

57. Dearden GI, Ravishankar V, Sakata K-T, Menon AK, Bergdoll L (2024) Protocol for the production and reconstitution of VDAC1 for functional assays. STAR Protoc 5:103240. 10.1016/j.xpro.2024.103240

58. Ando T, Kodera N, Takai E, Maruyama D, Saito K, Toda A (2001) A high-speed atomic force microscope for studying biological macromolecules. Proc Natl Acad Sci U S A 98:12468–12472. 10.1073/pnas.211400898

59. Schneider CA, Rasband WS, Eliceiri KW (2012) NIH Image to ImageJ: 25 years of image analysis. Nat Methods 9:671–675. 10.1038/nmeth.2089

60. Horcas I, Fernández R, Gómez-Rodríguez JM, Colchero J, Gómez-Herrero J, Baro AM (2007) WSXM: a software for scanning probe microscopy and a tool for nanotechnology. Rev Sci Instrum 78:013705. 10.1063/1.2432410

61. Ritchie TK, Grinkova YV, Bayburt TH, Denisov IG, Zolnerciks JK, Atkins WM, Sligar SG (2009) Chapter 11 - Reconstitution of membrane proteins in phospholipid bilayer nanodiscs. Methods Enzymol 464:211–231. 10.1016/S0076-6879(09)64011-8

62. Rostovtseva TK, Kazemi N, Weinrich M, Bezrukov SM (2006) Voltage gating of VDAC is regulated by nonlamellar lipids of mitochondrial membranes. J Biol Chem 281:37496–37506. 10.1074/jbc.M602548200

63. Narahari AK, Kreutzberger AJ, Gaete PS, Chiu Y-H, Leonhardt SA, Medina CB, Jin X, Oleniacz PW, Kiessling V, Barrett PQ, Ravichandran KS, Yeager M, Contreras JE, Tamm LK, Bayliss DA (2021) ATP and large signaling metabolites flux through caspase-activated Pannexin 1 channels. Elife 10:e64787. 10.7554/eLife.64787

64. Rostovtseva TK, Weinrich M, Jacobs D, Rosencrans WM, Bezrukov SM (2024) Dimeric Tubulin Modifies Mechanical Properties of Lipid Bilayer, as Probed Using Gramicidin A Channel. International Journal of Molecular Sciences 25:2204. 10.3390/ijms25042204

65. Queralt-Martín M, Bergdoll L, Jacobs D, Bezrukov SM, Abramson J, Rostovtseva TK (2019) Assessing the role of residue E73 and lipid headgroup charge in VDAC1 voltage gating. Biochim Biophys Acta Bioenerg 1860:22–29. 10.1016/j.bbabio.2018.11.001

66. Teijido O, Rappaport SM, Chamberlin A, Noskov SY, Aguilella VM, Rostovtseva TK, Bezrukov SM (2014) Acidification asymmetrically affects voltage-dependent anion channel implicating the involvement of salt bridges. J Biol Chem 289:23670–23682. 10.1074/jbc.M114.576314

67. Rappaport SM, Teijido O, Hoogerheide DP, Rostovtseva TK, Berezhkovskii AM, Bezrukov SM (2015) Conductance hysteresis in the voltage-dependent anion channel. Eur Biophys J 44:465–472. 10.1007/s00249-015-1049-2

68. Dodda LS, Cabeza de Vaca I, Tirado-Rives J, Jorgensen WL (2017) LigParGen web server: an automatic OPLS-AA parameter generator for organic ligands. Nucleic Acids Res 45:W331–W336. 10.1093/nar/gkx312

69. Dodda LS, Vilseck JZ, Tirado-Rives J, Jorgensen WL (2017) 1.14*CM1A-LBCC: Localized Bond-Charge Corrected CM1A Charges for Condensed-Phase Simulations. J Phys Chem B 121:3864–3870. 10.1021/acs.jpcb.7b00272

70. Jorgensen WL, Tirado-Rives J (2005) Potential energy functions for atomic-level simulations of water and organic and biomolecular systems. Proc Natl Acad Sci U S A 102:6665–6670. 10.1073/pnas.0408037102

71. Berendsen HJC, Postma JPM, van Gunsteren WF, Di Nola A, Haak JR (1984) Molecular dynamics with coupling to an external bath. Journal of Chemical Physics 81:3684–3690. 10.1063/1.448118

72. Bussi G, Donadio D, Parrinello M (2007) Canonical sampling through velocity rescaling. J Chem Phys 126:014101. 10.1063/1.2408420

73. Parrinello M, Rahman A (1981) Polymorphic transitions in single crystals: A new molecular dynamics method. Journal of applied physics 7182–7190

74. Darden T, York D, Pedersen L (1993) Particle mesh Ewald: An N⋅log(N) method for Ewald sums in large systems. Journal of Chemical Physics 10089–10092

75. Páll S, Hess B (2013) A flexible algorithm for calculating pair interactions on SIMD architectures. Computer Physics Communications 184:2641–2650. 10.1016/j.cpc.2013.06.003

76. Hess B, Bekker H, Berendsen HJC, Fraaije JGEM (1997) LINCS: A linear constraint solver for molecular simulations. Journal of Computational Chemistry 18:1463–1472. 10.1002/(SICI)1096-987X(199709)18:12<1463::AID-JCC4>3.0.CO;2-H

77. Souza PCT, Thallmair S, Conflitti P, Ramírez-Palacios C, Alessandri R, Raniolo S, Limongelli V, Marrink SJ (2020) Protein-ligand binding with the coarse-grained Martini model. Nat Commun 11:3714. 10.1038/s41467-020-17437-5

78. Gowers RJ, Linke M, Barnoud J, Reddy TJE, Melo MN, Seyler SL, Domanski J, Dotson DL, Buchoux S, Kenney IM, Beckstein O (2019) MDAnalysis: A Python Package for the Rapid Analysis of Molecular Dynamics Simulations. In: Report Number: LA-UR-19-29136. Research Org.: Los Alamos National Laboratory (LANL), Los Alamos, NM (United States)

79. Michaud-Agrawal N, Denning EJ, Woolf TB, Beckstein O (2011) MDAnalysis: a toolkit for the analysis of molecular dynamics simulations. J Comput Chem 32:2319–2327. 10.1002/jcc.21787

80. Işık M, Bergazin TD, Fox T, Rizzi A, Chodera JD, Mobley DL (2020) Assessing the accuracy of octanol–water partition coefficient predictions in the SAMPL6 Part II log P Challenge. J Comput Aided Mol Des 34:335–370. 10.1007/s10822-020-00295-0

81. Calculation of molecular lipophilicity: State-of-the-art and comparison of log P methods on more than 96,000 compounds - Mannhold - 2009 - Journal of Pharmaceutical Sciences - Wiley Online Library. https://onlinelibrary.wiley.com/doi/10.1002/jps.21494. Accessed 10 July 2026

82. Daina A, Michielin O, Zoete V (2017) SwissADME: a free web tool to evaluate pharmacokinetics, drug-likeness and medicinal chemistry friendliness of small molecules. Sci Rep 7:42717. 10.1038/srep42717

83. Daina A, Michielin O, Zoete V (2014) iLOGP: a simple, robust, and efficient description of n-octanol/water partition coefficient for drug design using the GB/SA approach. J Chem Inf Model 54:3284–3301. 10.1021/ci500467k

84. Cheng T, Zhao Y, Li X, Lin F, Xu Y, Zhang X, Li Y, Wang R, Lai L (2007) Computation of octanol-water partition coefficients by guiding an additive model with knowledge. J Chem Inf Model 47:2140–2148. 10.1021/ci700257y

85. Wildman SA, Crippen GM (1999) Prediction of Physicochemical Parameters by Atomic Contributions. Journal of Chemical Information and Computer Sciences 39:868–873. 10.1021/ci990307l

86. Lipinski CA, Lombardo F, Dominy BW, Feeney PJ (2001) Experimental and computational approaches to estimate solubility and permeability in drug discovery and development settings. Adv Drug Deliv Rev 46:3–26. 10.1016/s0169-409x(00)00129-0

87. Klimovich PV, Shirts MR, Mobley DL (2015) Guidelines for the analysis of free energy calculations. J Comput Aided Mol Des 29:397–411. 10.1007/s10822-015-9840-9

88. Andreasen MD, Souza PCT, Schiøtt B, Zuzic L (2024) Creating Coarse-Grained Systems with COBY: Towards Higher Accuracy in Membrane Complexity. bioRxiv. 10.1101/2024.07.23.604601

89. Bernetti M, Bussi G (2020) Pressure control using stochastic cell rescaling. J Chem Phys 153:114107. 10.1063/5.0020514

90. Kim H, Fábián B, Hummer G (2023) Neighbor List Artifacts in Molecular Dynamics Simulations. J Chem Theory Comput 19:8919–8929. 10.1021/acs.jctc.3c00777

91. Awasthi N, Hub JS (2019) Free-energy calculations of pore formation in lipid membranes. In: Biomembrane Simulations. CRC Press, pp 109–124

92. Kumar S, Rosenberg JM, Bouzida D, Swendsen RH, Kollman PA (1992) THE weighted histogram analysis method for free-energy calculations on biomolecules. I. The method. Journal of Computational Chemistry 13:1011–1021

93. Kroon PC, Grunewald F, Barnoud J, Tilburg M van, Brasnett C, Souza PCT de, Wassenaar TA, Marrink S-JJ (2025) Martinize2 and Vermouth: Unified Framework for Topology Generation. eLife 12:. 10.7554/eLife.90627.3

94. Böhm R, Amodeo GF, Murlidaran S, Chavali S, Wagner G, Winterhalter M, Brannigan G, Hiller S (2020) The Structural Basis for Low Conductance in the Membrane Protein VDAC upon β-NADH Binding and Voltage Gating. Structure 28:206–214.e4. 10.1016/j.str.2019.11.015

95. Ester M, Kriegel H-P, Sander J, Xu X (1996) A density-based algorithm for discovering clusters in large spatial databases with noise. In: Proceedings of the Second International Conference on Knowledge Discovery and Data Mining. AAAI Press, Portland, Oregon, pp 226–231

96. Pedregosa F, Varoquaux G, Gramfort A, Michel V, Thirion B, Grisel O, Blondel M, Müller A, Nothman J, Louppe G, Prettenhofer P, Weiss R, Dubourg V, Vanderplas J, Passos A, Cournapeau D, Brucher M, Perrot M, Duchesnay É (2018) Scikit-learn: Machine Learning in Python

