## Supplementary figures for "A Membrane-Disruptive Action of VBIT-4 Challenges Its Role as a Widely Used VDAC1 Oligomerization Inhibitor"

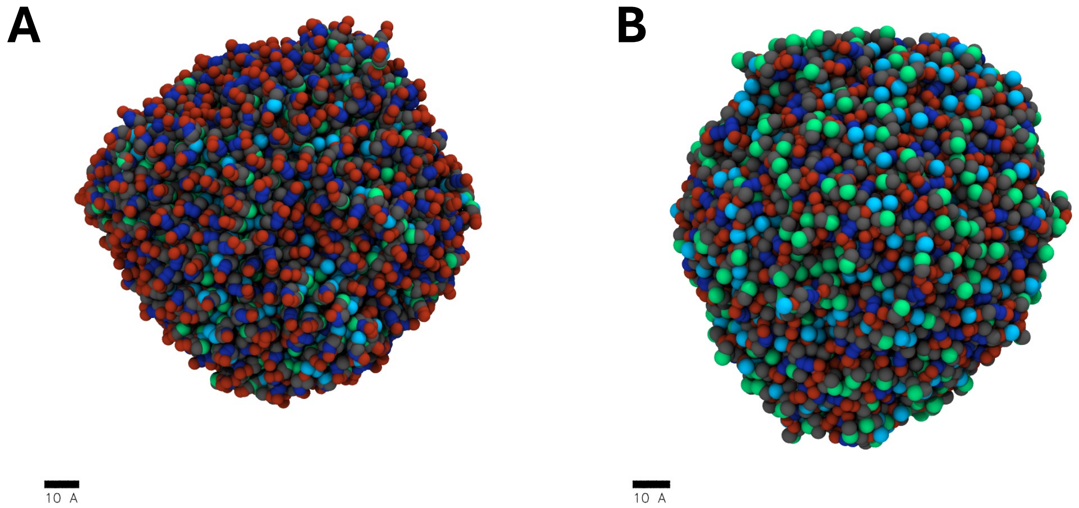


**Supplementary Figure 1. VBIT-4 self-assembles into micelle-like aggregates in aqueous solution.** Molecular dynamics simulations of 1000 VBIT-4 molecules under a charged (A) or neutral state (B) in a 15 × 15 × 15 nm³ water box show spontaneous clustering into compact micelle-like structures, consistent with the compound's amphiphilic and hydrophobic properties.

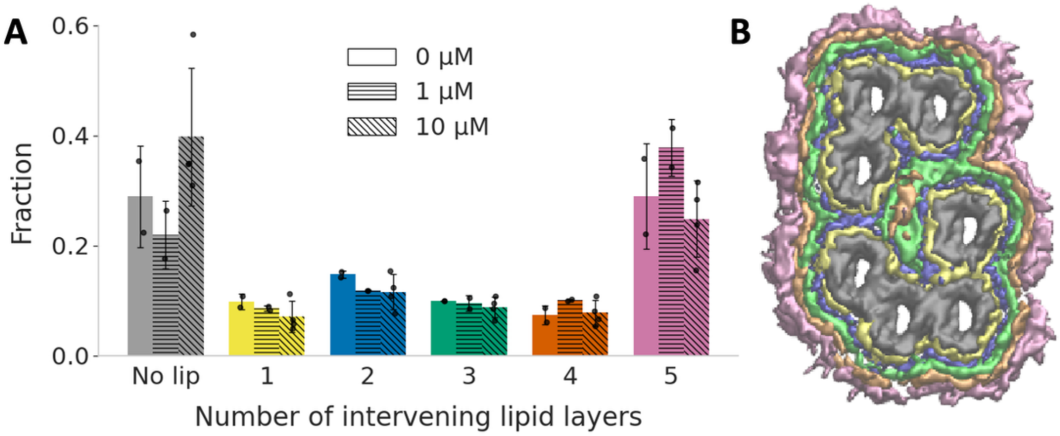


**Supplementary Figure 2. VBIT-4 does not alter the spatial organization of VDAC pairs in AFM micrographs.** (A) Distribution of inter-protein distances measured without VBIT-4 (open bars; 829 distances from 2 AFM fields), with 1 µM VBIT-4 (horizontal hatching; 380 distances from 2 fields), and with 10 µM VBIT-4 (oblique hatching; 568 distances from 4 fields). The number of intervening lipid layers between proteins were derived from distances between the proteins: 0 lipids (<46 Å, grey), 1 (46-50 Å, yellow), 2 (50-56 Å, blue), 3 (56-62 Å, green), 4 (62-68 Å, orange), and ≥5 (> 68 Å, purple). Error bars represent standard deviations calculated per AFM field. (B) Schematic illustrating a cluster of six proteins separated by various numbers of lipid layers (color-coded as in A). Inter-protein distances were determined using Delaunay triangulation of detected protein positions, which constructs a network of nearest-neighbor connections by linking points such that no additional point lies within the circumcircle of any triangle. This method restricts analysis to local neighbors, minimizing contributions from non-local associations and thereby providing accurate estimates of lipid-mediated spacing within clusters (total N = 5,331 distances across 8 images, each analyzed in triplicate).

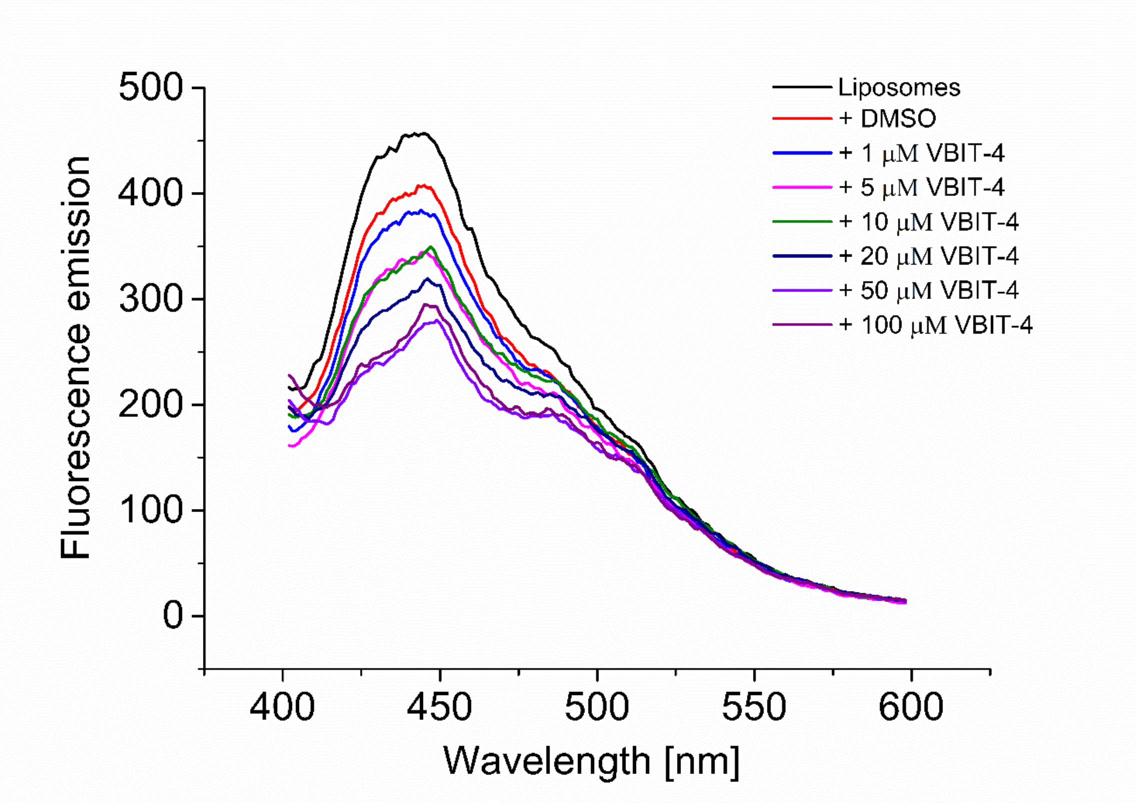


**Supplementary Figure 3. Laurdan fluorescence emission spectra** at different concentrations of VBIT-4 (excitation wavelength 375 nm). Laurdan fluorescence is sensitive to the polarity and hydration of the lipid bilayer, allowing detection of changes in lipid packing and water penetration within the membrane. Shifts in Laurdan's generalized polarization (GP), therefore, provide a quantitative measure of membrane perturbation. A clear VBIT-4-induced shift in

Laudran emission is visible, characteristic of a change in the environment around the dye. The DMSO concentration was constant for all the conditions.

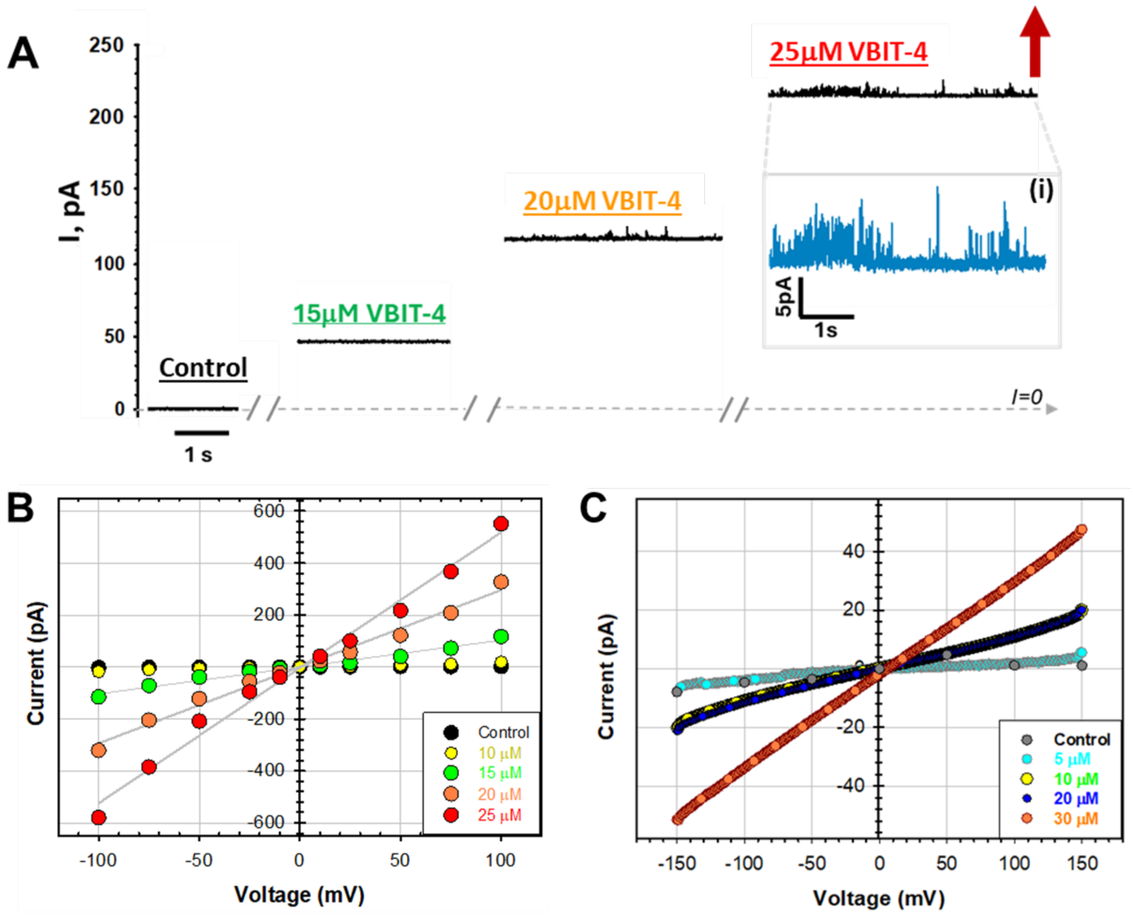


**Supplementary Figure 4. A.** Representative current records obtained on the PLM made from PLE before and after the addition of 15, 20, and 25 µM of VBIT-4 to both sides of the membrane at 50 mV applied voltage. Large fast fluctuations of the conductance at 25 µM of VBIT-4 (inset (*i*) shows the current trace at 25 µM VBIT-4 at a finer current scale) preceded the membrane rupture shown by the upward red arrow. Dashed grey lines indicate a zero current. The membrane bathing solutions consisted of 150 mM KCl buffered with 5 mM HEPES at pH 7.4. The current was digitally filtered using a 500 Hz Bessel (8-poles) filter for presentation. **B.** The current-voltage (I/V) curves obtained in experiments shown in (A) at different VBIT-4 concentrations. Grey lines are linear regressions indicating the Ohmic behavior of VBIT-4-induced conductances. **C.** To test whether the differences in membrane preparation account for the variation in reported VBIT-4 effects, we compared “dry” PLMs with “painted” membranes formed by Mueller’s method, which incorporates organic solvent (decane). Painted membranes were prepared using the Orbit Mini system (Nanion Technologies, Munich, Germany; see Methods). Shown are representative I/V curves from the same painted planar membrane composed of DPhPC in decane at increasing VBIT-4 concentrations added to the *cis* (grounded) compartment. The medium was 150 mM KCl and 5 mM HEPES (pH 7.4). Starting at 5 µM, VBIT-4 reproducibly induced leakage in both painted and "dry" membranes composed of DPhPC or PLE, confirming that its permeabilizing activity is independent of the bilayer formation method.

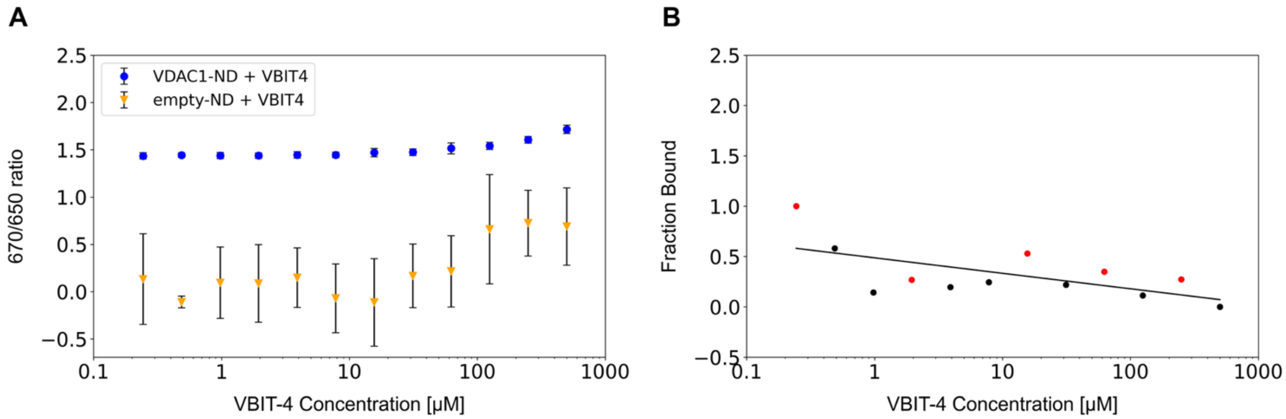


**Supplementary Figure 5.** Additional Microscale Thermophoresis (MST) data. **A.** Spectral shift at 670/650 nm of VDAC1-nanodiscs and lipid nanodiscs does not show any difference, indication of the absence of a specific binding site for VBIT-4 on VDAC1. **B.** Titration of VBIT-4 into labelled MSP1D1 without lipids. The MSP-only binding check showed no aggregations and a measurable fluorescent count. Upon addition of VBIT4, there was aggregation in capillaries 1, 4, 7, 9, 11 (data points marked in red). The response amplitude and SNR were too low to analyse (≈ 0.8). Data were plotted as fraction bound = $(F-F_{min})/(F_{max}-F_{min})$.

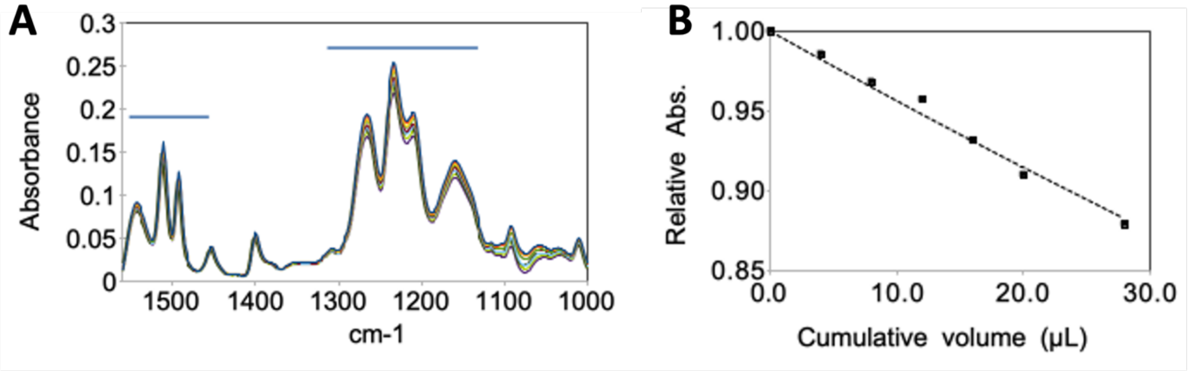


**Supplementary Figure 6.** Fourier-transform infrared spectroscopy (**FTIR) analysis reveals poor solubility and strong aggregation of VBIT-4.** To assess VBIT-4 solubility, we used FTIR, which detects molecular vibrations and distinguishes dissolved from aggregated species through characteristic spectral shifts—making it well suited for amphiphilic molecules like VBIT-4. Measurements were performed in attenuated total reflectance (ATR) mode, with the sample deposited directly on a diamond crystal serving as an infrared-transparent support. **A.** FTIR spectra of VBIT-4 showing its characteristic absorption bands at 1470–1570 $cm^{-1}$ and 1125–1300 $cm^{-1}$. From the upper to lower traces, six successive washings of the adsorbed material were performed. Despite repeated buffer exchanges (4 µL each), the spectra remained unchanged, indicating that VBIT-4 remained bound to the crystal surface, consistent with aggregation rather than solubilization. This behavior correlates with the turbid appearance of VBIT-4 suspensions in buffer, whereas clear concentrated solutions can be obtained by adding excess acid, even with moderate DMSO (20%). **B.** Relative absorbance of VBIT-4, computed by integrating the 1125–1300 $cm^{-1}$ region in (A) after successive washes with phosphate buffer. The dashed line represents a desorption model indicating that only ~0.44% of the adsorbed VBIT-4 was resolubilized per microliter of wash, confirming its strong aggregation and limited aqueous solubility.

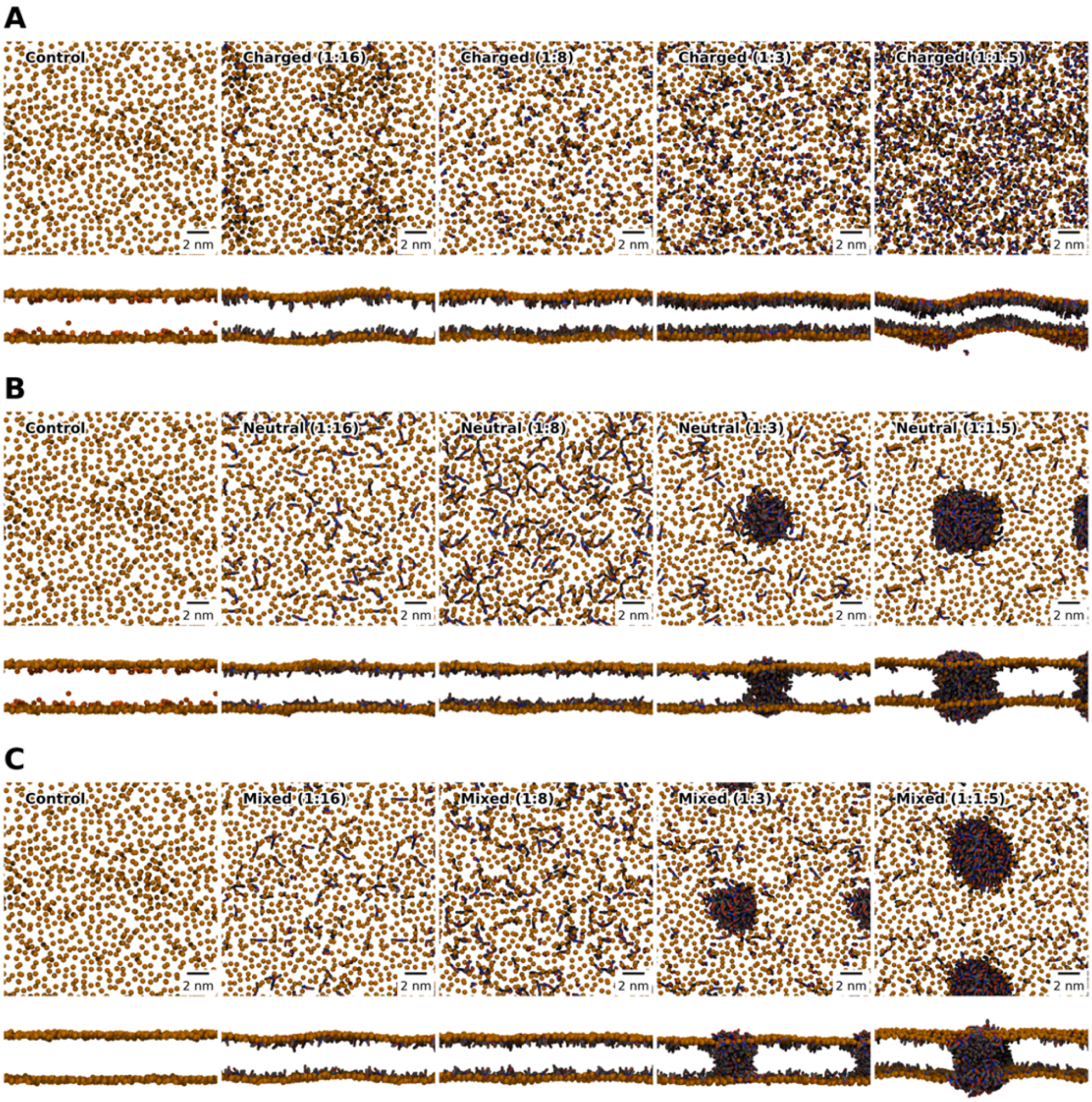


**Supplementary Figure 7. VBIT-4 forms pore-like structures through phase separation**. Representative coarse-grained MD snapshots showing the effect of increasing VBIT-4 concentrations on mitochondrial outer membrane mimics after 10 μs of simulation. Lipid phosphate beads are shown in orange, and VBIT-4 molecules in grey, with chemically distinct beads highlighted in red and blue. Top and side views are displayed for (A) charged, (B) neutral, and (C) 50:50 charged-to-neutral mixtures of VBIT-4 at increasing VBIT-4:Lipid ratios (0, 1:16, 1:8, 1:3, 1:1.5). Given that the pKa of the piperazine group is close to physiological pH, VBIT-4 is unlikely to exist exclusively in a charged or neutral form. Its protonation state likely depends on the local environment—being predominantly neutral when deeply inserted in the bilayer and charged near the phosphodiester/glycerol region. Consequently, pore formation at high concentrations likely involves both species: neutral VBIT-4 forming the membrane core, while charged molecules accumulate near the lipid headgroups or within the pore lumen, promoting water penetration and defect stabilization. To approximate this mixed state, simulations of a 50:50

charged-to-neutral mixture were performed, which showed aggregation and pore-like organization similar to neutral VBIT-4 but with intermediate membrane perturbation behavior.

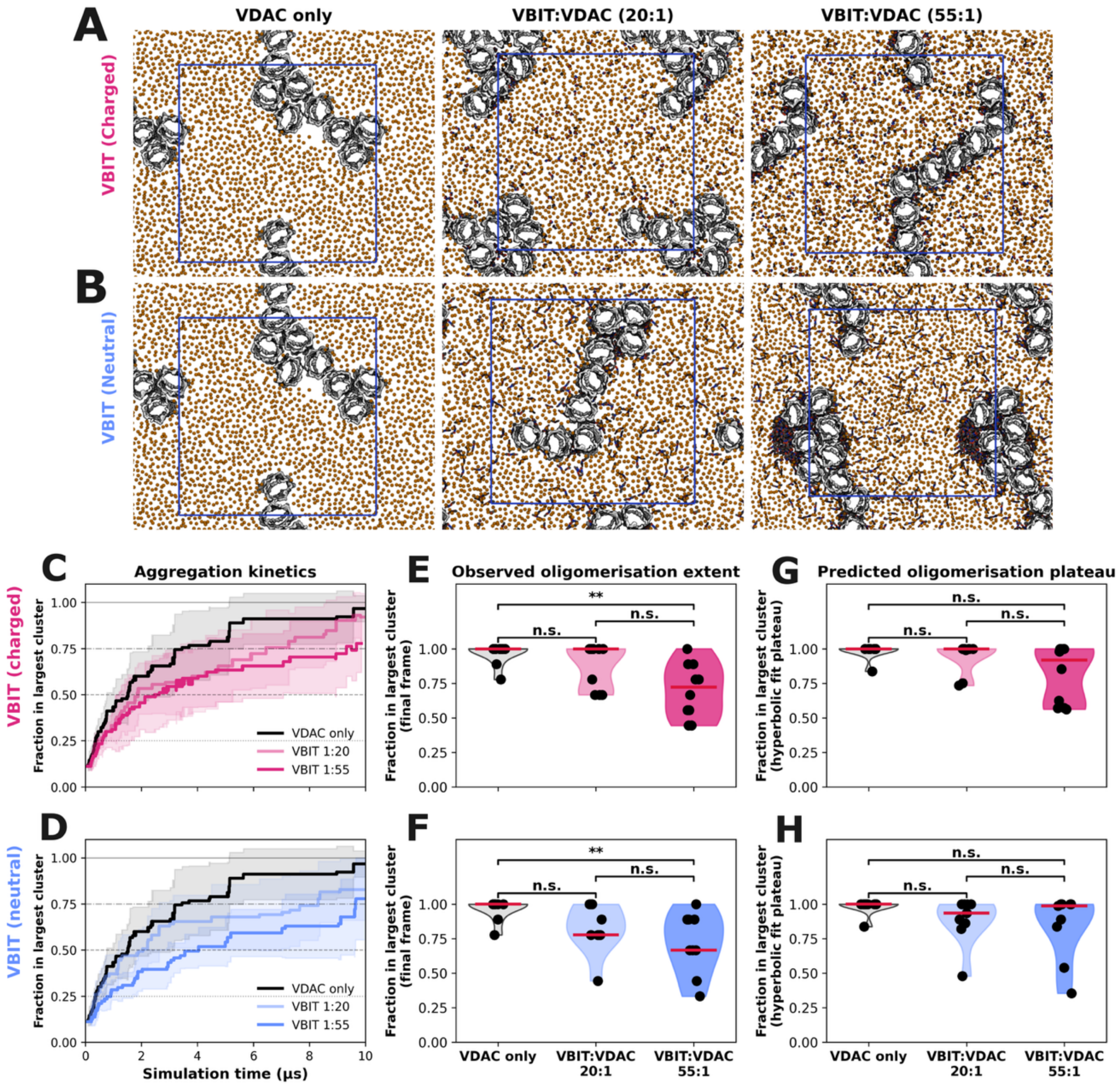


**Supplementary Figure 8. VBIT-4 does not prevent VDAC-1 oligomerisation, but slows it down.** Representative simulation snapshots at 10 μs for charged (**A**) and neutral (**B**) VBIT4 protonation states at increasing VBIT:VDAC1 ratios (left to right: VDAC only, 20:1, 55:1). VDAC1 proteins are shown in grey; lipid phosphate beads are shown in orange; VBIT-4 in grey, with chemically distinct beads highlighted in red and blue; the blue box indicates the periodic boundary unit cell. Mean aggregation kinetics (± SD, shaded) for charged (**C**) and neutral (**D**) VBIT-4 conditions. The fraction of proteins in the largest cluster is shown over 10 μs. The mean and standard deviation values are obtained from 10 independent simulations for each condition. Black: VDAC1 only; light/dark colour: VBIT-4 1:20 and 1:55, respectively. Observed oligomerisation extent at 10 μs (final frame value) for charged (**E**) and neutral (**F**) conditions. Predicted oligomerisation plateau estimated by hyperbolic curve fitting to each replicate trajectory, for charged (**G**) and neutral (**H**) conditions. Violins show the distribution across 10 replicates; the red line indicates the median. Statistical comparisons by Mann-Whitney U test with Holm correction; ** $p < 0.01$, n.s. not significant.

Notably, the right panel of B shows nucleation of VBIT-4–induced pores in proximity to VDAC1. This likely reflects local membrane perturbations associated with VDAC1, including thinning and lipid scrambling activity, which may facilitate VBIT-4 redistribution between leaflets and promote pore formation.

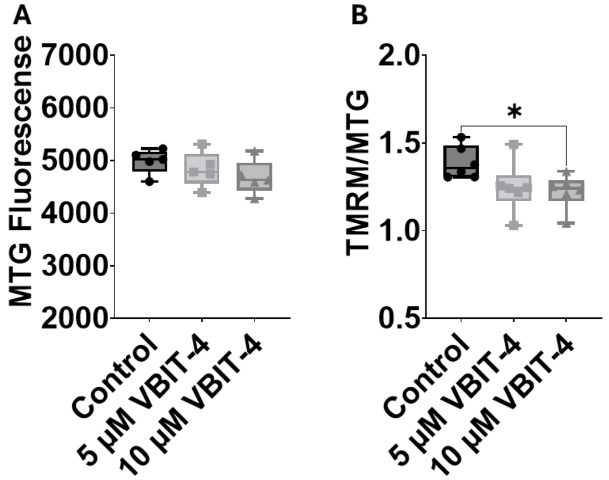


**Supplementary Figure 9. A.** Flow cytometry measurements of mitochondrial mass using mitotracker green (MTG) upon the addition of 5 µM (■) and 10 µM (▲) VBIT-4 compared to vehicle control. **B.** TMRM fluorescence normalized to MTG fluorescence (TMRM/MTG). Data from 6 independent experiments are represented in the box plots. The symbols represent data from independent experiments. The borders of the boxes define the 25th and 75th percentiles, with the median displayed as black lines and error bars indicating the standard deviation from the mean. Significance was tested using one-way ANOVA followed by the Dunnett post hoc test (*$p < 0.05$).

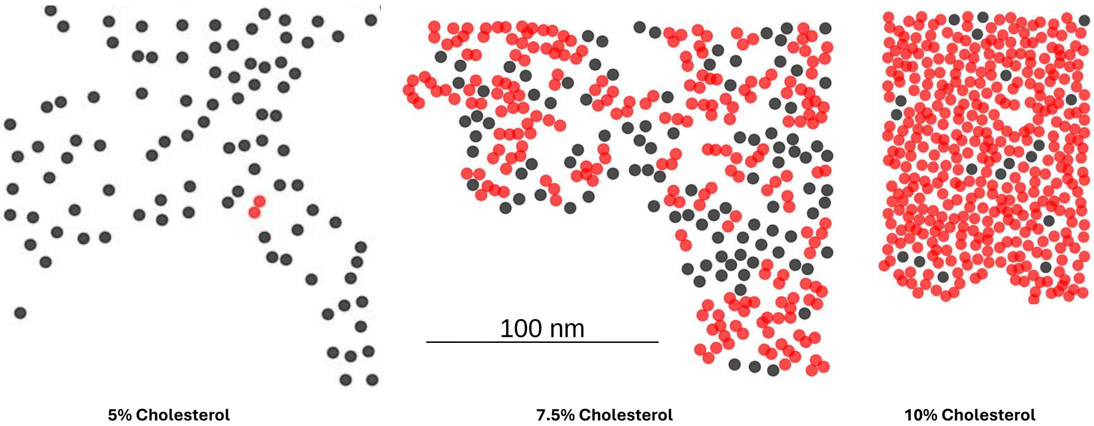


**Supplementary Figure 10. Changing the membrane composition affects the size and compaction of VDAC1 assemblies.** Figure adapted from Lafargue et al. [17]. This figure illustrates the cluster assignment for VDAC1 assemblies within a mitochondria outer membrane mimicking membrane, characterized by varying cholesterol concentrations, ranging from 5% to 10% cholesterol. VDACs forming direct contacts with another VDAC1 are represented in red. Isolated VDAC1 monomers are depicted in black. It is clearly evident that the number of VDAC1 with direct contacts with other VDACs, and available for cross-linking (all colour dots), are highly variable in the different conditions.

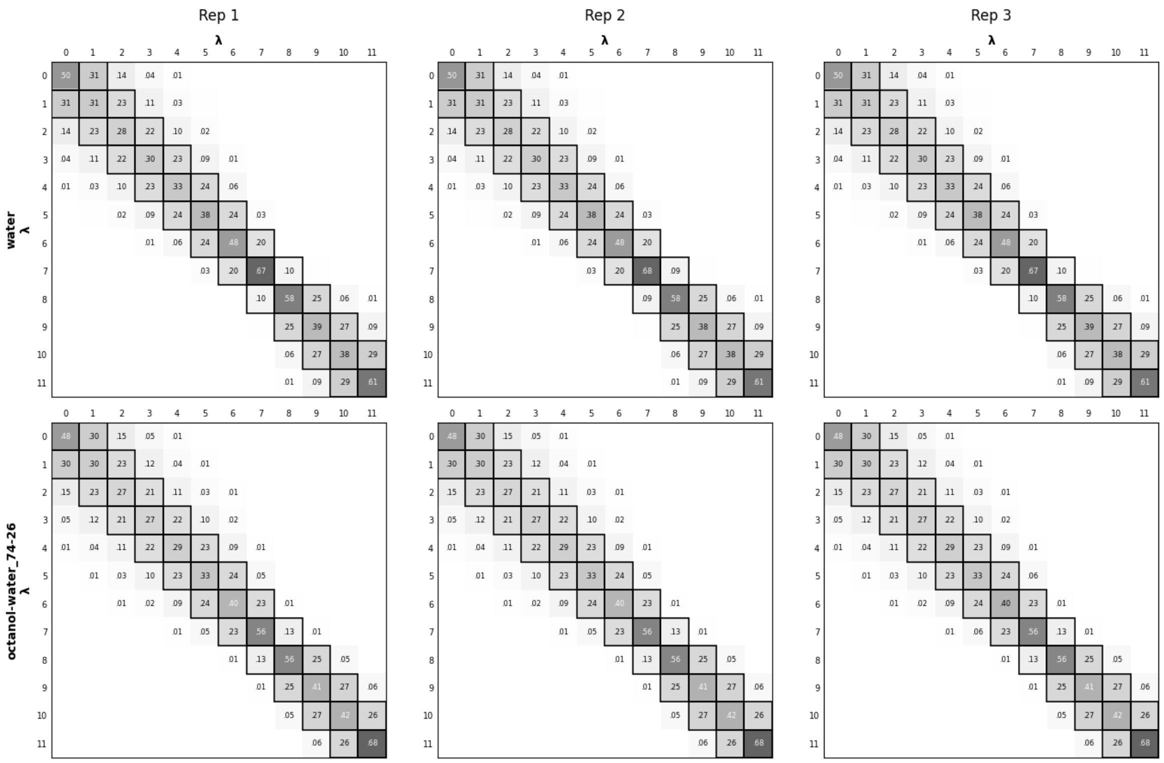


**Supplementary Figure 11.** MBAR overlap matrices for the water (top) and octanol-water 74:26 (bottom) alchemical legs across three independent replicates. Each cell reports the overlap between lambda windows *i* and *j*; values closer to 1 indicate greater phase-space overlap and more reliable free energy estimates between those states. Bold borders highlight the diagonal and first off-diagonal elements, representing the nearest-neighbour overlaps most critical for convergence. All adjacent lambda pairs exceed the minimal overlap threshold of 0.03, confirming adequate phase-space sampling across the full alchemical pathway.

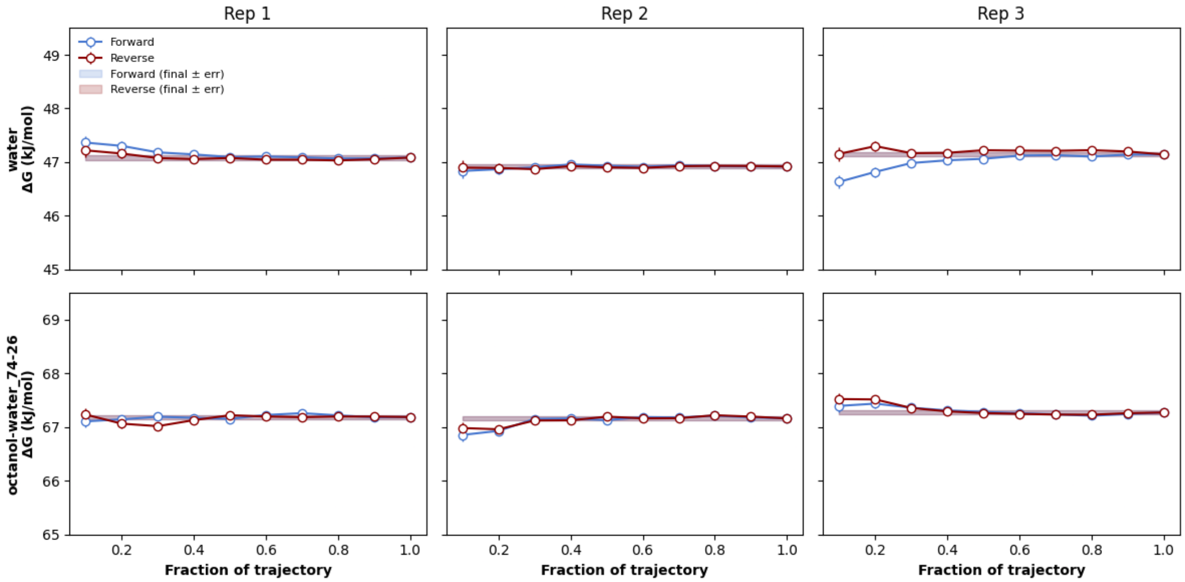


**Supplementary Figure 12.** Forward and reverse MBAR convergence of the solvation free energy for water (top) and octanol-water 74:26 (bottom) across three independent replicates. The forward estimate is computed using increasing fractions of the trajectory from the beginning, while the reverse estimate uses increasing fractions from the end. Rapid convergence of both estimates and their agreement within error at the full trajectory (shaded bands) indicates well-equilibrated simulations. All six legs show stable free energy estimates.

| PLM Lipid composition | KCl, M | Number of experiments | $G_m$, pS 10 µM VBIT-4 | $G_m$, pS 20 µM VBIT-4 | $G_m$, pS 50 µM VBIT-4 |
|---|---|---|---|---|---|
| **DPhPC** | 0.15 | 2 | 30; 230 | 540 | |
| **PLE** | 0.15 | 6 | 180; 590 | 3200 | 850; 1110 |
| **PLE** | 1 | 3 | 29; 200 | 316; 97 | 3260 |
| **PC:PE:chol** | 0.15 | 3 | 54; 45 | 670; 806 | 319; 3630 |

**Supplementary Table 1.** Summary of electrophysiology experiments performed on planar lipid membranes (PLM) of three lipid compositions – DPhPC, PLE, and DOPC:DOPE:chol (60:32.5:7.5) - in membrane bathing solutions of 0.15 or 1.0 M KCl, buffered with 5 mM HEPES at pH 7.4. The effect of VBIT-4 on membrane conductance ($G_m$) was measured ~15 min after the addition of VBIT-4 to one side (cis) of the membrane at an applied voltage of 100 mV. Each $G_m$ corresponds to the highest conductance measured at a given VBIT-4 concentration; those shown in the same color in each row correspond to the $G_m$ values measured on the same membrane. $G_m$ varies significantly across experiments, even under the same experimental conditions. Some PLMs ruptured upon the addition of 10 µM VBIT-4, while others remained stable until higher concentrations. Most membranes ruptured at ~30 µM, although a subset resisted up to 50 µM. Notably, 50 µM was the highest concentration at which PLMs remained intact for at least 15 min after VBIT-4 addition at an applied voltage of 100 mV. Only the PLMs with conductance < 1 pS in control before VBIT-4 addition were used in all experiments.
